# CRISPR activation screens map the genomic landscape of cancer glycome remodeling

**DOI:** 10.1101/2025.05.26.656133

**Authors:** John Daly, Lidia Piatnitca, Mohammed Al-Seragi, Vignesh Krishnamoorthy, Simon Wisnovsky

## Abstract

Many cancer types upregulate expression of sialic acid-containing glycans. These oligosaccharides subsequently engage inhibitory Siglec receptors on immune cells, allowing cancer cells to evade immune surveillance. The genetic mechanisms by which this glycome remodeling occurs remain poorly defined. Understanding the ways that cancer cells change their cell surface glycosylation is critical for identification of biomarkers and targets for glycan-directed immunotherapy. In this study, we performed multiple gain-of-function CRISPR activation (CRISPRa) screens to broadly define genetic pathways that regulate expression of Siglec-binding glycans. We show that Siglec ligand expression is largely controlled through genetic competition between genes that catalyze α2-3 sialylation and GlcNAcylation of galactose residues. Perturbation of enzyme expression at this key biosynthetic node provides multiple “paths” by which cancers can acquire elevated expression of Siglec ligands. We further show that cancer glycome remodeling is aided by overexpression of novel “professional ligands” that facilitate Siglec-glycan binding. Notably, we also find that expression of the CD24 gene is genetically dispensable for cell-surface binding of the inhibitory receptor Siglec-10. Finally, by integrating our functional genetic model with clinical tumor genomic data, we identify the sulfotransferase enzyme GAL3ST4 as a potential novel driver of immune evasion in glioma cells. Taken together, this study provides a first-in-class genomic atlas to aid understanding of cancer-associated glycosylation and identifies immediately actionable targets for cancer immunotherapy.

## Introduction

In 2018, the Nobel Prize in Medicine recognized the growing impact of immunotherapy as a strategy for treating cancer. Immunotherapies work by blocking the receptor-ligand interactions that restrain anticancer immunity. These drugs thus quench inhibitory signaling and re-activate potent anticancer immune responses^1–3^. Immunotherapies have produced remarkable results, even in patients with advanced cancer or otherwise untreatable disease^4,5^. However, many patients still fail to respond to existing immune therapies^6^. This is likely because different cancers exhibit significant heterogeneity in gene and cell-surface marker expression^7^. Immune cells also express dozens of inhibitory receptors that may be relevant in different patients and cancer subtypes^1–3^. So far, only a few receptor-ligand pairs have approved inhibitors^8^. Characterizing new mechanisms of immune suppression in cancer is thus crucial for the development of next-generation cancer therapeutics.

Changes in cell-surface glycosylation can play a critical role in suppressing the anticancer immune response^9^. Immune cells express several families of receptors that bind to glycan ligands. The Siglec (<u>si</u>alic acid-binding immuno<u>g</u>lobulin-like <u>lec</u>tin) family of receptors, for example, are characterized by their shared affinity for glycans that contain the sialic acid monosaccharide (Fig. 1A)^10–12^. Binding between a Siglec and its ligands triggers phosphorylation of ITIM/ITSM motifs located on the intracellular face of the receptor^10–12^. Subsequent recruitment of the inhibitory phosphatase SHP-1 to the cell membrane represses immune activation^10–12^. In recent years, it has been shown that a many cancers “remodel” their cell-surface glycome so as upregulate expression of glycan ligands for these Siglec receptors^13–17^. This process is sometimes termed *cancer-associated hypersialylation* and occurs in a remarkably wide cross-section of different tumor types^18,19^.

**Figure 1.**
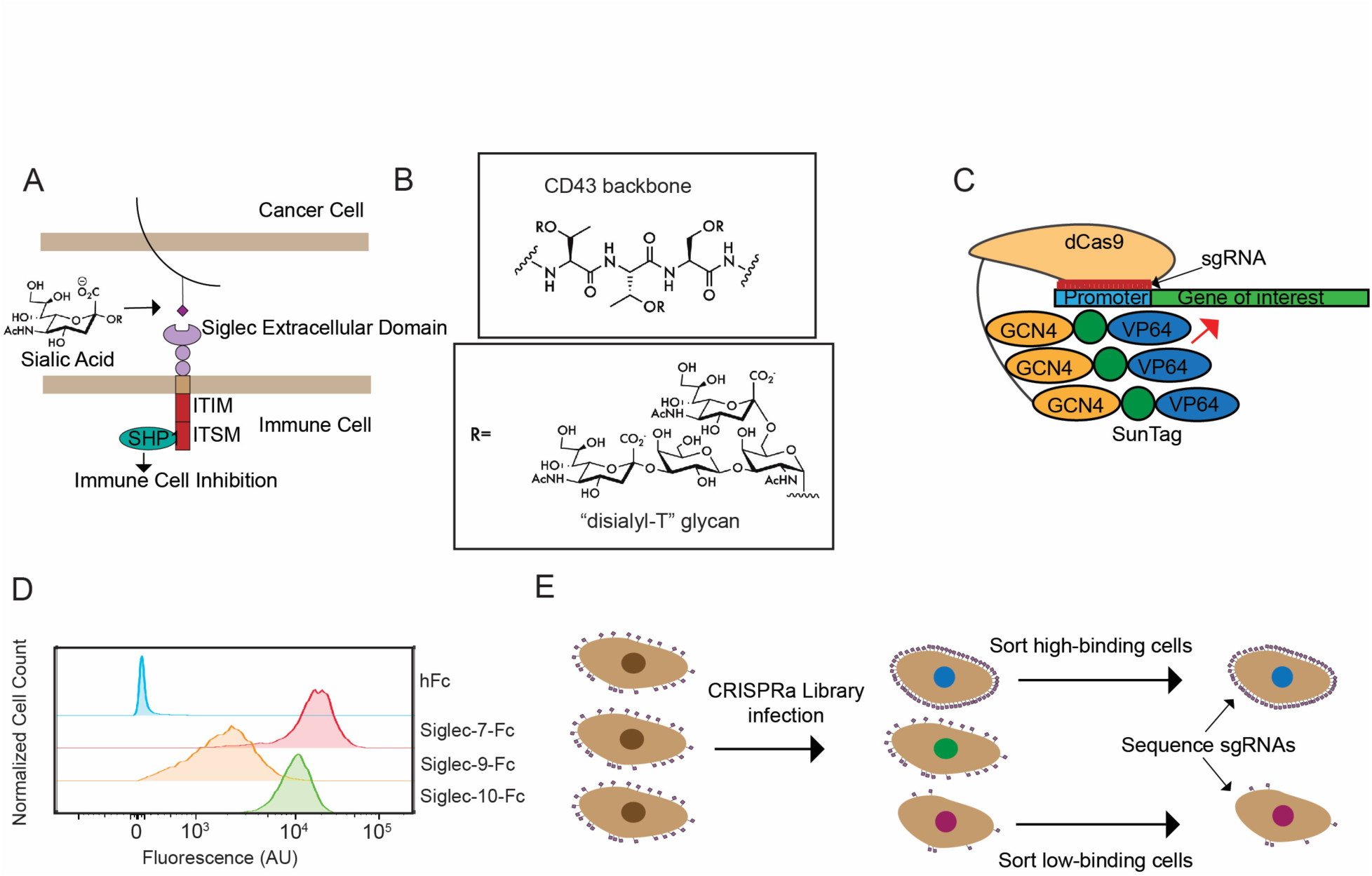
A CRISPR activation (CRISPRa) screening strategy for dissecting genetic drivers of Siglec ligand expression. **A)** Inhibition of anticancer immunity by Siglec receptors. Siglecs bind to sialic acid-containing glycans on the surface of target cells. **B)** Components of the Siglec-7 ligand structure. Disialyl-T glycans are organized in dense arrays on the backbone of specific mucin-type O-glycoproteins. **C**) Targeted overexpression of genes by CRISPRa. A cell line expressing a dCas9-GCN4 peptide array and a VP64-SunTag fusion protein serves as the target cell line (K-562-CRISPRa cells). This cell line is lentivirally transduced with an sgRNA targeting the promoter region of a target gene. Subsequent recruitment of chromatin remodeling factors induces an increase in transcriptional activity. **D)** Recombinant Siglec-Fc proteins were precomplexed with an AlexaFluor647-antihuFc antibody (1 μg/mL) for 1 hour on ice. Precomplexes were subsequently incubated with K-562 cells for 30 minutes and analyzed by flow cytometry. Representative flow cytometry plots are depicted for each Siglec-Fc as well as a human Fc (hFc) negative control. **E)** K-562-CRISPRa cells were lentivirally transduced with a genome-wide library of ∼104,000 sgRNAs (5 sgRNAs/gene). After selection and propagation, 1.25 x 10^8^ cells were then stained with Siglec-Fc reagents as in **B**. A high-binding (top 20%) and low-binding (bottom 20%) of the fluorescent population was then selected and sorted by FACS.

Increased biosynthesis of Siglec-binding glycans (Siglec ligands) is thus a common phenomenon in cancer. However, the specific genetic mechanisms that produce this phenotype remain poorly characterized. Sialic acid is incorporated into many distinct oligosaccharide chains on cell-surface glycoproteins and glycolipids^11^. Siglec-glycan interactions can also be influenced by many factors, including: 1) the monosaccharide composition and stereochemistry of the glycan, 2) secondary interactions with non-carbohydrate elements of protein scaffolds, and 3) the valency of glycan presentation on the cell surface^20–22^. Siglec ligand biosynthesis thus requires coordinated expression and/or repression of many genes encoding specific glycan-modifying enzymes and cell-surface scaffolds. Additionally, there are likely upstream oncogenes (transcription factors, kinases, etc.) that more broadly regulate glycosylation by controlling expression of multiple elements in these polygenetic circuits^14,23^. Systematic identification of these key genes could provide a wealth of new targets for the development of immunotherapeutic inhibitors.

Cancer glycome remodeling is thus an emergent phenotype with a highly complex genetic basis. In prior work, we began to interrogate this phenomenon using high-throughput functional genomic screening. In a pilot study, we applied a CRISPR interference (CRISPRi) screening strategy to produce an initial map of genes that are required for Siglec-7 ligand expression in hematopoietic cancer cells^24^. Cells were transduced with a genome-wide library of sgRNAs, stained with fluorescently labeled Siglec-7-Fc, and sorted by FACS to isolate cells with decreased binding.

Sequencing was then performed to identify sgRNAs that produce this loss of Siglec-7 ligand expression. This study yielded several novel insights. We found that Siglec-7-Fc binding required expression of a small cluster of genes that includes the sialyltransferases (STs) ST6GALNAC1 and ST3GAL2, the glycosyltransferase (GT) C1GALT1, and the cell-surface scaffolding protein CD43. Subsequent biochemical characterization revealed that Siglec-7 binds to an O-linked tetrasaccharide (the “disialyl-T” antigen) that is synthesized by these biosynthetic enzymes^24^ (Fig. 1B). We also showed that Siglec-7/disialyl-T binding is enhanced through clustering of adjacent glycans on specific mucin scaffolding proteins like CD43^24^ (Fig. 1B). Blocking or ablating expression of these glycans stimulated immune killing of leukemia in both cell and animal models^14,24,25^. This work thus defined a new target for leukemia immunotherapy and validated a general method to dissecting regulation of glycosylation in cancer^26^.

Loss-of-function (LOF) genetic screening can be a powerful approach to identifying regulators of cell-surface glycosylation. Indeed, several other studies have since applied LOF CRISPR screening to study a variety of other significant processes in glycobiology^27,28^. Other studies have also addressed some of these same questions using carefully designed, focused arrays of sgRNAs against glycan biosynthesis genes^29,30^. However, all these methods have significant methodological limitations. Firstly, cell-surface glycosylation patterns vary significantly in a cell and tissue-dependent manner. By some estimates, only ∼50% of glycan biosynthesis genes are even expressed in any given cell model^31,32^. A hit discovered in one LOF screen may thus have limited generalizability to other cell types or tissues. Secondly, LOF screening may not adequately assess functionally redundant or complementary genes. This is a particular problem for factors like GTs, which can have overlapping functions and substrate specificities^33^. Finally, genes that are essential to cell growth and viability can often be missed by traditional LOF screening. Our picture of how Siglec ligand biosynthesis is regulated in cancer cells thus remains incomplete and limited to a few select cell models.

We reasoned that CRISPR activation (CRISPRa) screening would be one ideal tool for addressing these problems. In CRISPRa screening, a cell line is engineered with a dCas9 protein that is tethered to a specific transcription activation domain (Fig. 1C)^34,35^. Subsequent transduction of cells with an sgRNA induces selective chromatin remodeling and targeted upregulation of gene transcription^34,35^. CRISPRa technology allows for large scale gain-of-function (GOF) genetic screens to be conducted in human cells^36^. CRISPRa screening has some key advantages over traditional CRISPR screening in the context of glycoscience research. Firstly, CRISPRa can transcriptionally activate many genes that have low baseline expression in a given cell line model^36–38^. By “forcing” overexpression of genes that would normally only be active in a specific tissue or developmental context, CRISPRa can assess a much broader range of possible regulators than LOF screening^36,37,39^. As a GOF screening technique, CRISPRa also eliminates the problems of gene essentiality and genetic redundancy that often arise in the study of glycan biosynthesis enzymes. In this study, we used CRISPRa screening technology to produce a genomic atlas of genes that regulate expression of ligands for Siglecs-7, -9 and -10. This first-in-class resource provides new, broadly applicable insights into the oncogenic mechanisms that drive cancer glycome remodeling across a range of tumor types. We subsequently leverage this dataset to characterize potential novel targets for cancer immunotherapy. Our key findings are outlined below.

## Results

### Genome-wide CRISPRa screens systematically map regulators of Siglec ligand expression

In previous work, we and others have shown that K-562 cells express ligands for some Siglec receptors^24,40^. K-562 cells are an ideal model for genome-wide screening, as they are genetically tractable and can be sorted by FACS in large numbers without loss of cell viability^24,26^. We therefore decided to use these cells as our model system for CRISPRa screening. K-562 cells were first transduced with constructs encoding dCas9-SunTag and VP64-scFv fusion proteins. This system allows for recruitment of multiple VP64 transcriptional activators to specific promoter sequences, thus upregulating expression of target genes^34^. We then incubated these cells (hereafter termed *K-562-CRISPRa* cells) with a panel of Siglec-Fc chimera proteins precomplexed with a fluorescent secondary antibody. For this experiment we chose to assess Siglecs-7, -9, -10 and -15, as all these receptors have recently been implicated as potential targets for immune checkpoint blockade in various tumor types^15,17,24,41,42^. Siglecs-7, -9 and -10 displayed significant binding to K-562s, confirming that cell-surface glycans are broadly hypersialylated in this cell line as observed previously (Fig. 1D)^24^. Siglec-15, conversely, showed no significant binding over background (Supplementary Fig. 1). We therefore chose to focus our subsequent studies on Siglecs-7, -9, and -10.

We transduced these K-562-CRISPRa cells with a genome-wide library of sgRNAs (∼104,000 sgRNAs, 5 sgRNAs/gene)^43^. Following selection, this library was stained with fluorescently labeled Siglec-7-Fc, Siglec-9-Fc or Siglec-10-Fc. Using FACS, we then sorted cells into either “low-staining” or “high-staining” bins. These populations corresponded respectively to the bottom 20% and the top 20% of the fluorescence distribution for each Siglec-Fc. Finally, we used next generation sequencing to quantitate the expression of sgRNAs in each of these populations. Using CasTLE^44^, we identified specific sgRNAs that were more abundant in either the low-staining or in the high-staining population. This allowed us to generate a list of gene hits whose overexpression either increases Siglec-Fc binding or decreases Siglec-Fc binding.

The full results of all three CRISPRa screens are plotted in Figs. 2A-C. CasTLE computes two distinct values for each gene: a CasTLE Score and an effect size. The CasTLE score is a measure of statistical significance based on the consistency of enrichment effects for different sgRNAs targeting the same gene. A higher CasTLE score indicates a greater level of statistical significance for a given hit. The effect size reflects the average difference in abundance of sgRNAs in the high and low staining populations. A higher effect size would indicate that gene upregulation exerts a greater impact on Siglec ligand expression. In our dataset, a positive effect size indicates that gene overexpression increased Siglec-Fc binding. These genes will subsequently be termed *positive regulators*. A negative effect size, conversely, suggests that gene overexpression causes a decrease in binding (*negative regulators*).

**Figure 2.**
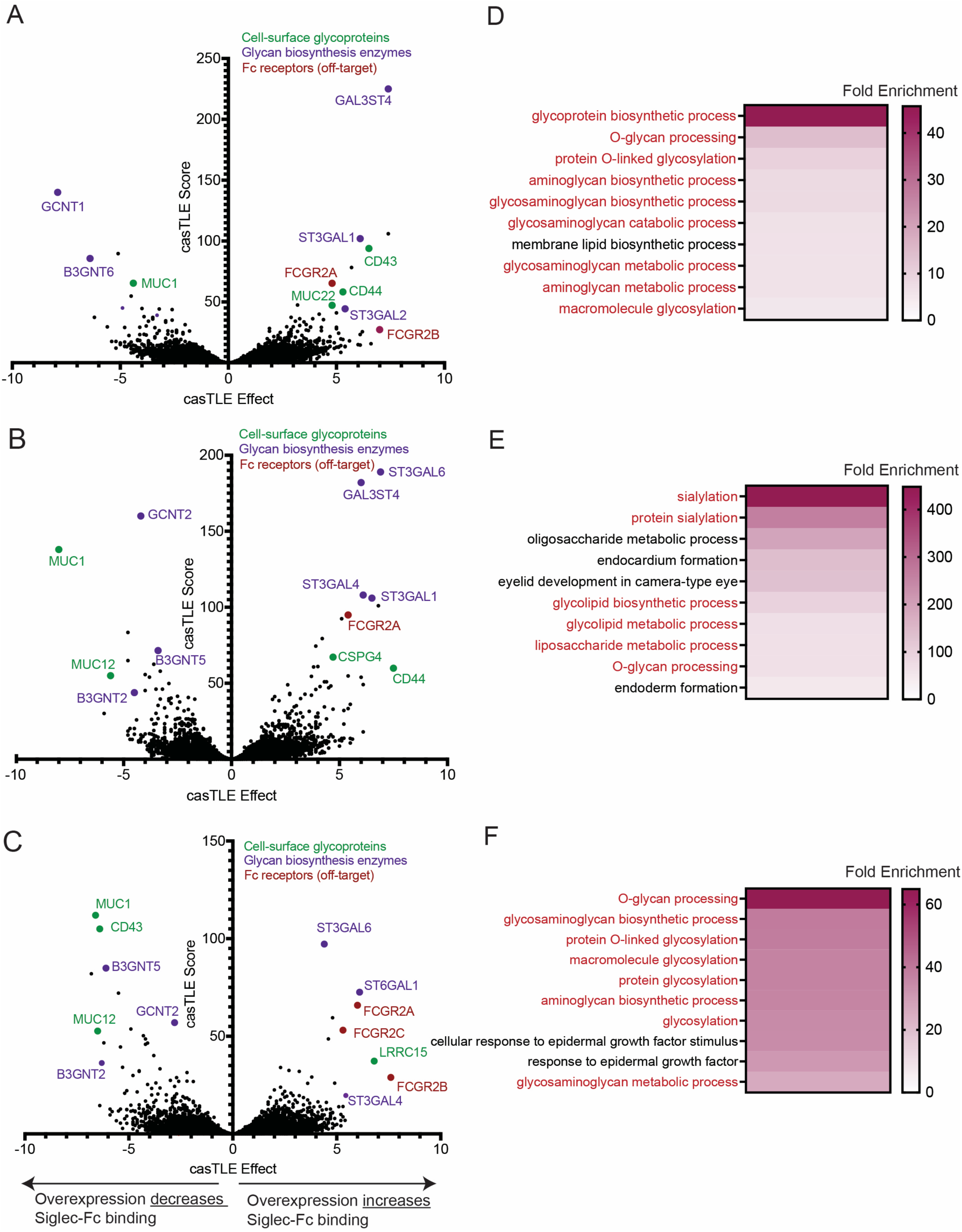
CRISPR activation screens reveal drivers of Siglec ligand expression. **A)**. CasTLE analysis was then performed to identify and rank hit genes from the Siglec-7-Fc screen. “CasTLE effect size” approximates the increase (or decrease) in the abundance of sgRNAs targeting a certain gene in the high-staining fraction relative to the low-staining fraction. A positive sign indicates gene overexpression increases Siglec-Fc binding, a negative sign indicates that gene overexpression decreases Siglec-Fc binding. “CasTLE Score” is a measure of statistical reproducibility of enrichment effects across multiple sgRNAs targeting the same gene. A higher CasTLE score indicates greater statistical confidence. All genes in library are plotted. Selected cell surface glycoproteins (green), glycan biosynthesis genes (purple) and Fc receptors (red) are highlighted on each graph. **B)** Analysis was performed as in **2A** for cells stained with Siglec-9-Fc. **C)** Analysis was performed as in **2A** for cells stained with Siglec-10-Fc. **D)** Gene Ontology (GO) enrichment analysis was performed using GOrilla. The top 10 terms by fold enrichment are ranked. All enrichments were statistically significant with an FDR (q-value) below 0.01. GO terms associated with carbohydrate biology and/or processing are highlighted in red. **E)** Analysis was performed as in **2D** for gene hits from the Siglec-9-Fc screen. **F)** Analysis was performed as in **2D** for gene hits from the Siglec-10-Fc screen.

We recovered many hits whose overexpression altered Siglec ligand expression. Several broad patterns were immediately apparent in the data. Firstly, GO term analysis of gene hits revealed heavy enrichment of functional terms related to glycan biosynthesis (Fig. 2D-F). Genes encoding sialyltransferase (ST) enzymes, which add sialic acid to glycans, were particularly prominent among the positive regulators identified in all three screens. We also identified several cell-surface proteins that directly bind to Siglec receptors. The gene encoding CD43 (SPN), for example, was the top positive regulator discovered in our Siglec-7 CRISPRa screen. These results broadly validate the success of our new screening method. We present our full, annotated results in Supplementary Table 1, in the hope that it will be a broadly useful resource for the glyco-immunology research community. We also highlight that over 50 % of our most significant hits (CasTLE score > 30) are natively expressed at extremely low levels in K-562 cells (Human Protein Atlas, mRNA TPM < 10) (Supplementary Table 2). This result confirms our hypothesis that CRISPRa can be used to identify tissue-specific regulators of Siglec ligand expression that may be missed by LOF screens. This “genomic atlas” thus provides a significantly broader, more generalizable view how cancer sialylation is regulated than any prior study. Below, we outline a set of the most important insights we have drawn and validated from these datasets.

### Siglec ligand expression is controlled by biosynthetic competition between sialylation and terminal branching/elongation of N- and O-linked glycans

We began by examining gene hits that are known to play a role in glycan biosynthesis. These were identified by association of genes with GO terms related to glycan metabolism (e.g., GO0005975, carbohydrate metabolic process). Unsurprisingly, many of these biosynthetic enzymes were STs. Figure 3A depicts the CasTLE scores for all ST enzymes encoded in the human genome. We observed that transcriptional upregulation of the ST3GAL family of STs produced particularly significant increases in Siglec ligand expression^45,46^. These enzymes catalyze the attachment of sialic acid onto the 3’ hydroxyl group of galactose residues^45,46^. This result accords with prior findings, as we and others have found that upregulation of ST3GAL enzymes drives elevated Siglec ligand biosynthesis in multiple cancer types^13,17,47^. The significance of ST3GAL genes in our screening dataset further confirms that galactose sialylation acts is a key “rate-limiting” step controlling expression of cell-surface Siglec ligands.

**Figure 3.**
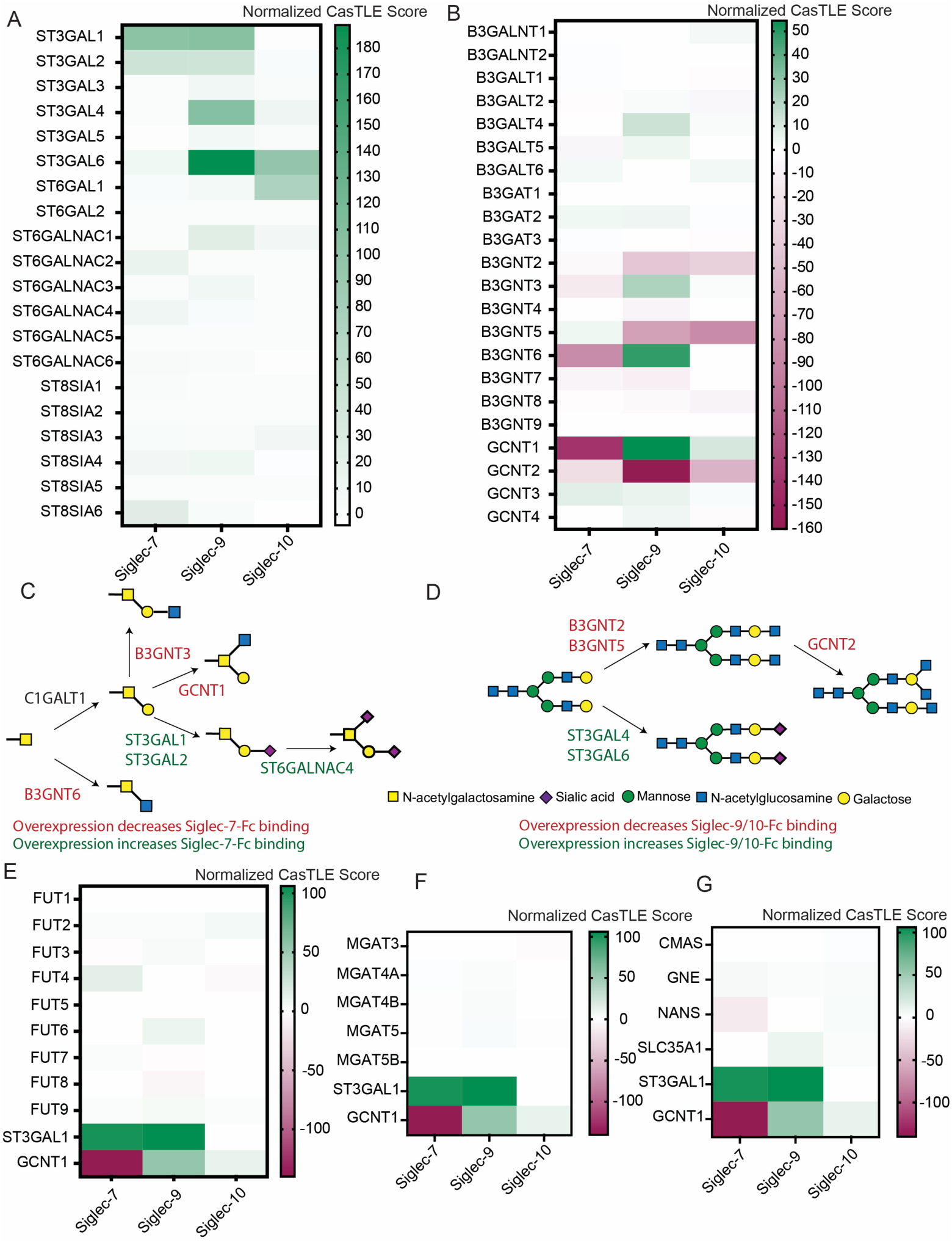
Siglec ligand expression is controlled by the transcriptional balance between enzymes involved in branching and sialylation. **A**) CasTLE scores for all known sialyltransferase genes in the human genome across three different CRISPRa screens. Scores have been given a sign to indicate directionality of effect. A positive score indicates that gene overexpression increased Siglec-Fc binding, a negative score indicates decreased binding. **B)** CasTLE scores for all genes encoding enzymes with β-Gal or β-GlcNAc transferase activity towards Galactose residues. Scores normalized as in **A**. **C)** Pathway for synthesis of O-linked glycans starting from αGalNAc. Biosynthetic enzymes that catalyze the relevant steps are highlighted. Genes with a CasTLE score > 30 in the Siglec-7 screen are highlighted in green, those with a CasTLE score < -30 are highlighted in red. **D)** Pathway for synthesis of N-linked glycans starting from an asialylated biantennary N-linked glycan. Biosynthetic enzymes that catalyze the relevant steps are highlighted. Genes with a CasTLE score > 30 in the Siglec-9/Siglec-10 screen are highlighted in green, those with a CasTLE score < -30 are highlighted in red. **E)** CasTLE scores for all genes encoding enzymes with fucosyltransferase activity. **F)** CasTLE scores for all genes encoding enzymes with GlcNAc-transferase activity towards Man residues. **G)** CasTLE scores for all genes encoding enzymes involved in sialic acid biosynthesis. ST3GAL1 and GCNT1 are shown as positive controls to normalize effect sizes.

We also found that overexpression of different ST3GAL family members produced distinct effects on the binding of different Siglec-Fc reagents. For example, Siglec-7 binding was increased by overexpression of both ST3GAL1 and ST3GAL2, but not any other enzymes in the family. Siglec-9 ligand expression was more plastic, with overexpression of all ST3GAL enzymes except ST3GAL5 having a significant positive effect. Finally, Siglec-10 ligand expression was increased only by upregulation of ST3GAL4 and ST3GAL6. ST6GAL1, which attaches sialic acid to the 6’ hydroxyl of galactose, was also a hit for Siglec-10. ST6GAL1 overexpression did not affect binding of Siglec-7-Fc or Siglec-9-Fc. These findings elegantly demonstrate that these different Siglecs bind distinct sialylated structures on the cell surface. ST3GAL1 and ST3GAL2, for example, are the primary STs that act on O-linked glycans^45,46^. These data thus support an emerging consensus that Siglec-7 primarily binds sialylated O-linked glycans on the surface of cells^14,17,24,29,48,49^. Conversely, the prominence of N-glycan active enzymes like ST3GAL4 and ST3GAL6 in our Siglec-9/10 screens demonstrates that these receptors likely bind N-linked glycans^13,17^. Cancers with distinct genetic alterations in ST3GAL genes are thus likely to exhibit major differences in cell-surface Siglec ligand expression, a finding with important implications for proper targeting of blocking therapeutics.

Interestingly, we recovered few ST hits outside of the ST3GAL family. ST6GALNAC4, which modifies the sialyl-T antigen to generate disialyl-T, was a weak but statistically significant hit in our Siglec-7 screen^14,50^. This result matches prior reports showing that this enzyme generates Siglec-7 ligands^14,29,51^. The enzyme ST8SIA6, which further modifies the disialyl T epitope to generate “trisialyl-T” antigens, was also recovered as a weak hit^52^. This finding accords with another recent study showing that the trisialyl T glycans can bind Siglec-7 with high affinity^52^. Apart from these genes, there were no other ST family enzymes that were recovered as statistically significant hits. For example, overexpression of STs involved in lipid sialylation (e.g, ST3GAL5, ST8SIA1) produced no effects on Siglec-Fc binding^53,54^. Siglec-Fc reagents often bind glycolipid-associated structures when they are presented on synthetic glycan arrays *in vitro*^10,11,55^. This type of binding, however, may not actually occur on an intact cell membrane. One explanation for this could be that glycolipids are quite small structures^56^. In an intact cell, these moieties are likely “buried” beneath a layer of N-linked and O-linked glycans that extend further out from the cell membrane^56^. Glycolipids may thus be largely inaccessible for physiological binding by Siglec receptors *in trans*, even if they do bind to Siglecs in some *in vitro* systems. If correct, these results indicate that modulating glycolipid sialylation is likely not a relevant mechanism by which cancer cells increase Siglec receptor engagement.

GTs from several other families were also recovered as hits. Figure 3B displays the CasTLE scores for all genes from the GCNT and B3GNT families, which append N-acetylglucosamine (GlcNAc) to various underlying glycan structures^57,58^. For these genes, the most notable effects were cases where gene overexpression significantly reduced Siglec-Fc binding. For example, upregulation of the enzymes GCNT1 and B3GNT3 was found to significantly antagonize Siglec-7 ligand expression. Given what is known about the substrate specificity of these enzymes, it is likely that that such genes modify core O-glycan structures such that they can no longer be acted on by STs (Fig. 3C)^51,59^. Overexpression of enzymes involved in GlcNAcylation of N-linked glycans (B3GNT2, B3GNT5 and GCNT2) produced a similar dampening effect on Siglec-9 and Siglec-10 ligand expression (Fig. 3D)^60^. Many of these enzymes act on the exact same underlying substrates as the ST3GAL enzymes we identified as strong positive regulators (Fig. 3C-D). “Biosynthetic competition” between ST genes and GlcNAcylation genes is thus clearly a major factor determining the overall density of Siglec ligands on the cell surface. Transcriptional dysregulation of genes at these key biosynthetic nodes is likely a major mechanism underlying cancer glycome remodeling^61^.

Finally, we noted several major classes of glycan-active enzyme that were not at all represented in our list of hits (Fig. 3E-G). These included enzymes involved in fucosylation (the FUT family), N-glycan branching (MGAT4/5), and biosynthesis of sialic acid monosaccharide precursors (CMAS, GNE, etc.)^57,62^. Several caveats must be noted here. Firstly, pooled genomic screening always suffers from false negatives due to variable sgRNA potency. Secondly, some biosynthetic steps may also already “saturated” due to already-high expression of a specific biosynthetic enzyme in our model cell line. For example, we did not obtain ST6GALNAC1 as a strong hit, despite having shown in previous LOF studies that knockdown of this gene reduces Siglec-7 biosynthesis. This effect is likely due to the expression of ST6GALNAC1 already being so high in K-562s as to make further overexpression of genes in this pathway ineffective^24^. Finally, biosynthesis of some glycan structures may require overexpression of multiple pathway components simultaneously, which is currently not possible at a genome-wide scale. With these limitations in mind, our results do imply that fucosylation, core N-glycan branching, and upregulation of CMP-sialic acid synthesis are all dispensable for achieving Siglec ligand overexpression in cancer cells. Instead, our data argue that Siglec ligand biosynthesis is mostly controlled by genetic competition between enzymes involved in sialylation and terminal branching/elongation of N- and O-linked glycans.

### Upregulation of distinct “professional ligands” enhances Siglec-7 and Siglec-9 binding

Some recent studies have suggested that Siglecs can exhibit preferential binding to “professional ligands”. Broadly defined, these are specific proteins that present glycans in an orientation or density that facilitates Siglec binding^22,24,41^. Overexpression of genes encoding these proteins may also be a mechanism underlying cancer glycome remodeling. We reasoned that our CRISPRa screening dataset might identify some of these factors. To test this hypothesis, we filtered our CRISPRa screen results for plasma membrane-localized genes that contained an extracellular domain (UniProt). We then used a stringent cut-off (CasTLE score > 30, effect size > 3) to identify “strong” hit genes that showed significant positive selection in at least one screen. These top hits are displayed in Figure 4A.

**Figure 4.**
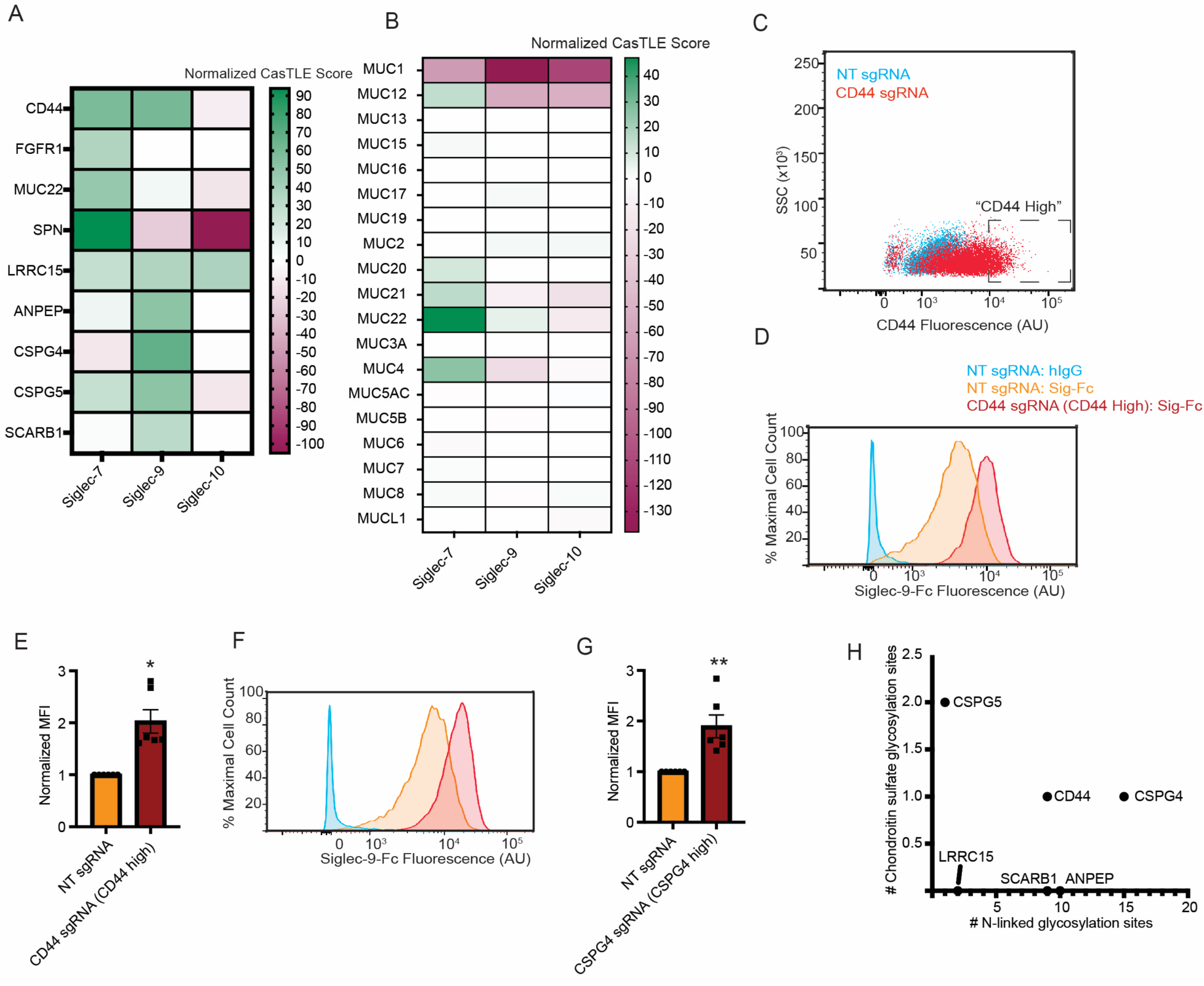
Overexpression of specific cell-surface glycoproteins increases binding of Siglec-7 and Siglec-9. **A**) CasTLE scores for all genes that had a CasTLE score > 30 in at least one screen and which contain an annotated extracellular domain (UniProt). CasTLE scores for all three screens are shown. **B)** CasTLE scores for all genes in the MUC family of mucin-type O-glycoproteins. **C)** K-562-CRISPRa cells were transduced with either a non-targeting sgRNA or an sgRNA against CD44. Cells were then stained with a fluorescent antibody against CD44 and analyzed by flow cytometry. The indicated gate (“CD44 High”) contains a population of cells with significantly elevated CD44 expression. **D)** K-562-CRISPRa cells were transduced as in **4C** and co-stained with an antibody against CD44 and Siglec-9-Fc. A representative flow cytometry plot is shown comparing Siglec-9-Fc staining in cells transduced with a non-targeting (NT) sgRNA to cells within the “CD44 High” expression gate. **E)** The median fluorescence intensity (MFI) of Siglec-9 staining is shown. MFI is internally normalized to the WT cell line. **F)** K-562 CRISPRa cells were transduced with an sgRNA against CSPG4 and co-stained with Sig9-Fc and a CSPG4 antibody as in **4C-D**. A representative flow cytometry plot is shown comparing Siglec-9-Fc staining in cells transduced with a non-targeting (NT) sgRNA to cells within the “CSPG4 High” expression gate. **G)** The MFI of Siglec-9 staining in NT cells vs. “CSPG4 High” cells is shown. MFI is internally normalized to the WT cell line. **H)** Top hits from the Siglec-9-Fc screen that contained an extracellular domain were analyzed in UniProt for the presence of different types of glycosylation sites. The number of annotated N-linked glycosylation sites for each gene is indicated on the x-axis, while the number of chondroitin sulfate modification sites is graphed on the y-axis. Statistical significance was determined using a two-tailed t-test. ** indicates p<0.01, * indicates p<0.05. Mean values plotted, error bars indicate SEM.

Unsurprisingly, many densely O-glycosylated proteins (mucins) were recovered as hits in our Siglec-7 screen. Our top hit was the SPN gene. This gene encodes CD43, a mucin we and others have shown interacts strongly with Siglec-7^24,61,63,64^. Of the five strong hits identified in our analysis, 4 genes (CD44, SPN, MUC22 and LRRC15) were found to contain mucin domains in a recent proteome-wide analysis^65^. The fifth gene (FGFR1) is a signaling receptor and so may regulate Siglec-7 ligand expression through indirect effects^66^. This trend was also confirmed by closer analysis of gene families that are known to encode mucin proteins. Several MUC genes (MUC12, MUC20, MUC21 and MUC4), for example, displayed significant positive effect sizes in our Siglec-7 screen (Fig. 4B). This effect was not observed for all mucins, however. Indeed, overexpression of the classical mucin MUC1 actually decreased Siglec-7-Fc binding^67^. These findings are consistent with prior studies, which have shown that only specific mucins can present disialyl-T glycans at a sufficient density to mediate Siglec-7 engagement^24,29^.

Once again, our CRISPRa screening strategy was clearly able to identify hit genes that are not natively expressed in K-562s. Such genes may thus be functionally relevant across a broader cross section of different cancer types. MUC4, for instance, is frequently overexpressed in pancreatic cancer and have been implicated as a key driver of pancreatic cancer tumorigenesis^68,69^. Given recent studies showing that Siglec-7 mediates immune suppression in pancreatic cancer, our results provide strong motivation for examining this specific protein as an immunotherapeutic target^16,17^. Similarly, other screen hits like MUC21 have been implicated as adverse prognostic markers in diseases like glioblastoma and melanoma^70^. Our dataset is thus a useful resource for characterization of novel Siglec-7 ligands in carcinomas, where known ligands like CD43 are unlikely to be expressed^61^.

The results of our Siglec-9 screen displayed a markedly different pattern. Here, there was no tendency for O-glycosylated proteins to be enriched as screen hits. Indeed, overexpression of many of the MUC genes, as well as SPN, decreased binding of Siglec-9-Fc (Fig. 4A-B). However, we did identify a distinct set of hits that are known to localize at the cell surface. Overall, 6 genes met our cut-off for statistical significance: LRRC15, CD44, CSPG4, CSPG5, ANPEP and SCARB1. These genes have never been described as Siglec-9 ligands. We therefore sought to confirm that overexpression of these genes increases Siglec-9-Fc binding.

We transduced K-562-CRISPRa cells with sgRNAs against the most significant three of these six top hit genes: CD44, CSPG4, and LRRC15. We then stained cells with fluorescent antibodies against each of these targets. In general, sgRNA transduction resulted in a marked increase in average protein expression, although with considerable intracellular heterogeneity (Fig. 4C). Because of this breadth in staining intensity, we next chose to co-stain cells with Siglec-Fc reagents and with protein-targeted fluorescent antibodies. This step allowed us to selectively analyze Siglec-Fc binding to cells that showed a significant increase in expression of the given cell-surface protein (Fig. 4C). We found that overexpression of CD44 and CSPG4 significantly enhanced binding of Siglec-9-Fc (Fig. 4D-G). For LRRC15, we found that sgRNA transduction increased non-specific binding to our hFc control, indicating that this hit is likely non-specific (Supplementary Fig. 2). CSPG4 and CD44 may thus represent novel professional ligands for Siglec-9.

It is unclear what shared feature is possessed by CSPG4 and CD44 that makes them bind to Siglec-9-Fc so strongly. We and others have previously shown that Siglec-9-Fc binding depends primarily on expression of complex N-linked glycans^13,29,71^. Unsurprisingly, both proteins have several N-linked glycosylation sites. Additionally, we noticed that three of our five valid hits (excluding LRRC15) possess annotated chondroitin sulfate (CS) modification sites (Fig. 4H). This result is notable, as CS is a rare type of glycosylation^72^. At this point, it is not clear whether CS may play a role in facilitating Siglec-9 binding. It is unlikely that Siglec-9 is directly binding to CS. However, CS modification may cause these proteins to adopt a conformation that exposes N-linked glycans for Siglec-9 binding. Regardless, the identification of these three proteins as Siglec-9 ligands provides a strong starting point for further investigation into Siglec-9’s protein binding selectivity^72^. Like our novel Siglec-7 ligands, several of these genes have also been implicated as drivers of tumor progression in several cancer subtypes^73,74^.

### Siglec-10 does not selectively bind CD24, but displays broad affinity for multiple classes of N-linked sialoglycans

One of the most surprising aspects of our dataset was a negative result: the absence of the cell-surface protein CD24 as a hit in our Siglec-10 screen. Multiple prior studies have reported that Siglec-10 binds selectively to CD24^41,75^. We thus expected that overexpression of this gene would enhance Siglec-10-Fc binding. As CRISPRa screens are prone to false negatives, we confirmed this result using a targeted assay. We selected two sgRNAs against the CD24 gene from our genome-wide library, transduced them into K-562-CRISPRa cells, and stained cells with an anti-CD24 fluorescent antibody. Both sgRNAs produced significant upregulation of cell-surface CD24 expression (Fig. 5A). This result highlights the general efficiency of gene upregulation induced by sgRNAs in our genome-wide library.

**Figure 5.**
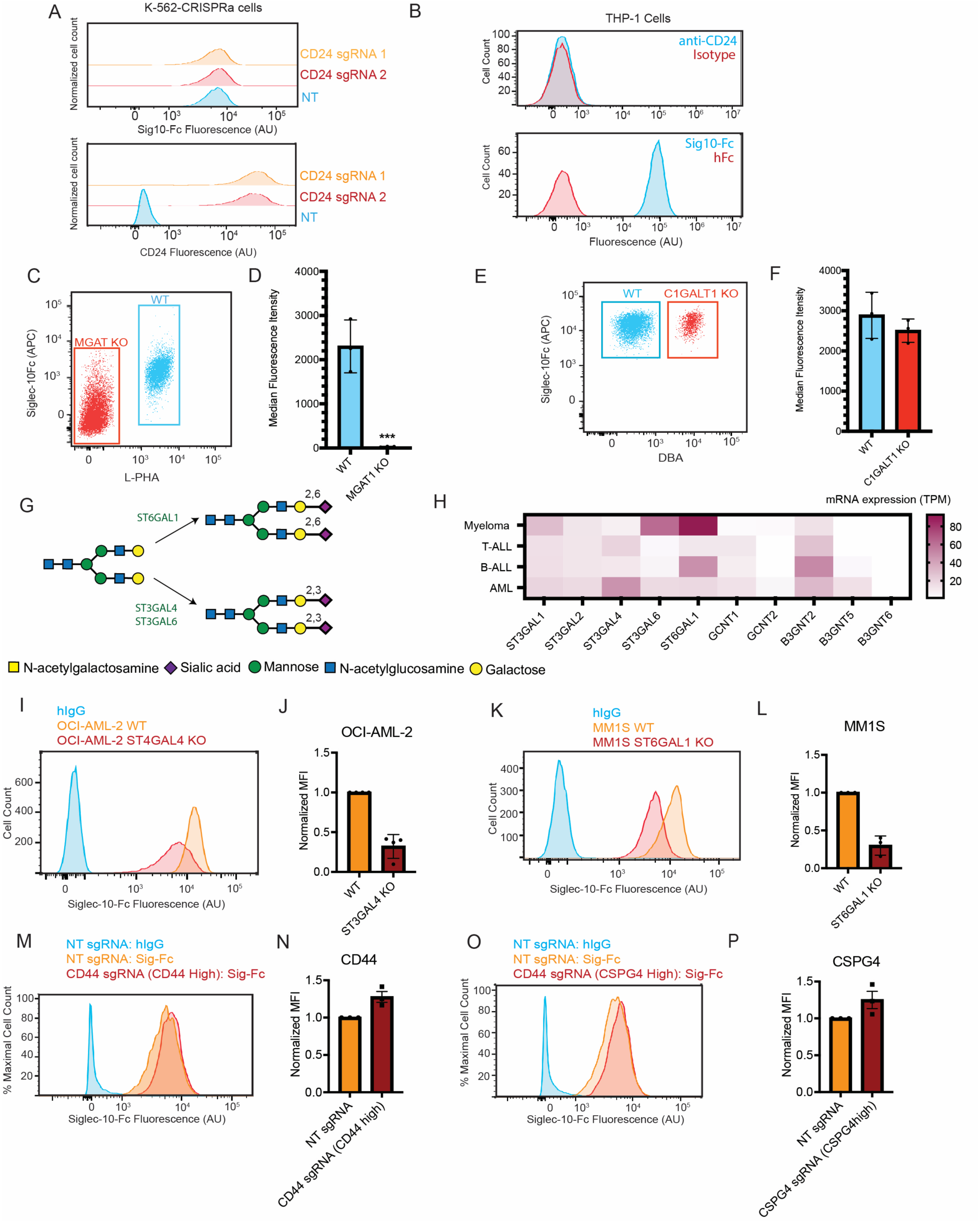
Siglec-10 displays broad specificity for N-linked sialoglycans. **A)** K-562-dCas9-CRISPRa cells were transduced with two different sgRNAs targeting the CD24 promoter region. After selection with puromycin, cells were stained with Siglec-10-Fc as in **1B** and with a fluorescent antibody against CD24. Representative flow cytometry plots for both stains are shown. **B)** THP-1 cells were stained with Siglec-10-Fc as in **1B** and a fluorescent antibody against CD24. Representative flow cytometry plots for both stains are shown. **C)** OCI-AML-2-Cas9 cells were lentivirally transduced with an sgRNA targeting MGAT1 to generate a polyclonal cell population containing WT and MGAT1 KO cells. Cells were then co-stained with Siglec-10-Fc and the lectin L-PHA (5 μg/mL), which binds to complex N-linked glycans. Representative staining is indicated on the flow cytometry plot. **D)** Graph indicates MFI of Siglec-10-Fc staining in the WT (L-PHA+) and MGAT1 KO (L-PHA-) populations. **E)** OCI-AML-2-Cas9 cells were lentivirally transduced with an sgRNA targeting C1GALT1. Cells were co-stained with Siglec-10-Fc and the lectin DBA, which binds to exposed α-GalNAc. Representative staining is indicated on the flow cytometry plot. **F)** Graph indicates MFI of Siglec-10-Fc staining in the WT (DBA-) and C1GALT1 KO (DBA+) populations. **G)** Pathway diagram indicates the enzymes involved in 2,3-linked and 2,6-linked sialylation of N-linked glycans. ST3GAL4, ST3GAL6 and ST6GAL1 were all strong hits in the Siglec-10 screen. **H)** The heat map indicates the average mRNA expression of key glycosyltransferase hits in cell lines derived from B-ALL, T-ALL, AML and multiple myeloma. mRNA expression values were extracted from DepMap and the Human Protein Atlas. **I)** OCI-AML-2-Cas9 cells were transduced an sgRNAs against ST3GAL4. Cells were then stained with fluorescently labeled Siglec-10-Fc. A representative flow cytometry plot is shown. **J)** The MFI of Siglec-10 staining in OCI-AML-2 WT and ST3GAL4 KO cells is indicated. MFI is internally normalized to the WT cell line. **K)** MM1S-Cas9 cells were transduced an sgRNAs against ST6GAL1. ST6GAL1 KO cells were isolated by FACS using the lectin SNA. Cells were then stained with fluorescently labeled Siglec-10-Fc. A representative flow cytometry plot is shown. **L)** The MFI of Siglec-10 staining in MM1S WT and ST6GAL1 KO cells is indicated. MFI is internally normalized to the WT cell line. **M)** K-562-CRISPRa cells were transduced and stained as in **4C**. A representative flow cytometry plot is shown comparing Siglec-10-Fc staining in cells transduced with a non-targeting (NT) sgRNA to cells within the “CD44 High” expression gate. **N)** The MFI of Siglec-10-Fc staining in NT cells vs. “CD44 High” cells is shown. MFI is internally normalized to the WT cell line. **O)** K-562 CRISPRa cells were transduced with an sgRNA against CSPG4 and co-stained with Sig10-Fc and a CSPG4 antibody as in **4C**. A representative flow cytometry plot is shown comparing Siglec-10-Fc staining in cells transduced with a non-targeting (NT) sgRNA to cells within the “CSPG4 High” expression gate. **P)** The MFI of Siglec-10 staining in NT cells vs. “CSPG4 High” cells is shown. MFI is internally normalized to the WT cell line. Statistical significance was determined using a two-tailed t-test. ** indicates p<0.01, * indicates p<0.05. Mean values plotted, error bars indicate SEM.

We then assessed whether CD24 overexpression was associated with increased Siglec-10-Fc binding. Surprisingly, non-transduced K-562-CRISPRa cells bound strongly to Siglec-10-Fc but did not express CD24 at detectable levels. CD24 overexpression also produced no change in Siglec-10-Fc binding (Fig. 5A). To further confirm this result, we stained another hematopoietic cell line (THP-1) with anti-CD24 and Siglec-10-Fc. These cells also expressed Siglec-10 ligands, but did not show any binding to our CD24 antibody (Fig. 5B). Our results in no way exclude the possibility that Siglec-10 may bind CD24 in some contexts. However, they do show that CD24 gene expression is not strictly necessary for Siglec-10 to bind the cell surface. Other ligands that modulate Siglec-10 signaling are thus likely to exist.

We next characterised the general structure of these Siglec-10 ligands. As mentioned above, the pattern of hits in our Siglec-10 screen strongly implied that Siglec-10 binds N-linked glycans. To confirm this result, we transduced OCI-AML-2 cells with Cas9 and an sgRNA against MGAT1, an essential enzyme in the synthesis of complex N-linked glycans^76^. OCI-AML-2 cells were chosen for these experiments because they strongly express ligands for both Siglec-9 and Siglec-10^13^. Cells were then co-stained with L-PHA, a plant lectin that binds N-linked glycans, and Siglec-10-Fc^76^. MGAT1 KO cells were identified by gating on the cell population that exhibited reduced L-PHA staining (Fig. 5C). These MGAT1 KO cells displayed a complete ablation of Siglec-10 ligand expression (Fig. 5D). In parallel, we also transduced cells with an sgRNA against C1GALT1, a key enzyme in elongation of O-linked glycans^76^. Here, C1GALT1 KO cells were identified by co-staining with the lectin DBA, which binds truncated O-glycans (Fig. 5E)^76^. C1GALT1 KO caused no change in Siglec-10-Fc staining (Fig. 5F). Together, these data confirm that Siglec-10 displays high selectivity for binding N-linked sialoglycans.

We next interrogated what types of N-linked sialoglycans are bound by Siglec-10. As mentioned above, our screen identified multiple STs that act on N-linked glycans. Two particularly strong hits were ST3GAL4 and ST3GAL6, which add sialic acid to terminal Galβ1,4-GlcNAc structures to generate Neu5Acα2,3-Galβ1,4-GlcNAc^45,77^. Interestingly, the other ST that showed a strong positive effect was ST6GAL1. This enzyme also acts on Galβ1,4-GlcNAc to generate Neu5Acα2,6-Galβ1,4-GlcNAc (Fig. 5G). ST6GAL1 only emerged as a hit in our Siglec-10 screen (Fig. 3A). These results imply that Siglec-10 can bind N-linked glycans that terminate with either a 2,3-linked or 2,6-linked sialic acid.

To confirm this result, we next assessed how CRISPR-Cas9 KO of these STs affects Siglec-10 ligand expression. Relative expression of ST3GAL4, ST3GAL6 and ST6GAL1 varies significantly across different cell and tissue types. To help select appropriate models for these experiments, we analyzed mRNA expression of these genes in over 100 blood cancer cell lines (Human Protein Atlas). As we have previously described, acute myeloid leukemia (AML) cell lines showed elevated expression of the gene ST3GAL4 relative to other genes in the ST family (Fig. 5H)^13^. We thus hypothesized that ST3GAL4 is likely the key driver of Siglec-10 ligand expression in AML. We transduced a representative AML cell line (OCI-AML-2) with Cas9 and an sgRNA against ST3GAL4. We confirmed successful gene knockout (over 90% editing) using TIDE analysis^13,78^ (Supplementary Fig. 3). Knockout of ST3GAL4 significantly reduced Siglec-10-Fc binding, demonstrating that Siglec-10 can indeed bind to N-linked glycans containing Neu5Acα2,3-Galβ1,4-GlcNAc (Fig. 5I-J).

In contrast to AML, we found that multiple myeloma (MM) cell lines expressed much higher levels of ST6GAL1 than ST3GAL4 (Fig. 5H). Hypersialylation has also been implicated as a driver of immune evasion in MM^48^. Here, we reasoned that ST6GAL1 was likely to be the key ST involved in Siglec-10 ligand biosynthesis. We therefore transduced a representative MM cell line (MM1S) with an sgRNA targeting ST6GAL1. Cells were then stained with the lectin SNA, which binds 2,6-linked sialic acids, and sorted via FACS to yield a purified ST6GAL1 KO population (Supplementary Fig. 4)^76,79^. ST6GAL1 KO induced a strong decrease in Siglec-10-Fc binding (Fig. 5K-L). These results confirm that Siglec-10 can bind a broad range of N-linked sialoglycans. They also highlight that different STs are likely to drive overexpression of Siglec-10 ligands in different cancer types.

Lastly, we assessed whether Siglec-10 binds any distinct professional ligands other than CD24. Surprisingly, we were not able to identify a single cell-surface protein hit in our Siglec-10 screen. The only gene that passed our statistical cutoff was LRRC15, which we had previously shown is a non-specific hit whose overexpression increases hFc binding (Supplementary Fig. 2). To confirm this result, we used CRISPRa to overexpress the two genes (CSPG4 and CD44) that we had already identified as professional Siglec-9 ligands. In contrast to our results with Siglec-9, we found that overexpression of these three genes had little effect on Siglec-10-Fc binding (Fig. 5M-P). Again, CRISPRa screening will always produce some false negatives. It is completely possible that there are cryptic Siglec-10 ligands that have not been revealed by this specific dataset. However, these results suggest that Siglec-10 exhibits quite broad affinity for many N-linked glycoproteins, such that overexpression of any given protein has a minimal effect on binding. This aspect of Siglec-10’s biology makes it distinct from both Siglec-7 and Siglec-9, where increased binding can clearly be mediated by overexpression of genes encoding distinct cell-surface ligands.

### The sulfotransferase enzyme GAL3ST4 drives Siglec ligand expression in glioma cells

Finally, we developed an unbiased hit prioritization pipeline to identify any high-value cancer immunotherapy targets that may have been missed by our previous analyses. As described above, our CRISPRa screen had identified dozens of genes whose overexpression increases cell-surface binding of inhibitory Siglec receptors. We hypothesized that if one of these genes was a significant driver of immune evasion in a specific cancer subtype, then higher expression of that gene would likely be associated with poor patient survival in that cancer. We thus cross-referenced our list of top positive regulators (CasTLE Score > 30) with a recent study that used the Cancer Genome Atlas (TCGA) to identify adverse prognostic factors across 33 different cancer types^80^. We first assessed whether elevated mRNA expression of our positive regulators was significantly associated with worse patient survival. The details of our analysis are fully described in *Materials & Methods.* The full results are provided in Supplementary Table 3.

This step revealed several interesting insights. First, high expression of our top ST hits was not strongly associated with poor patient survival in any cancer type. Secondly, several cell-surface ligands (e.g, CD44 in renal cell carcinoma) were identified as key adverse prognostic markers. Finally, we identified a remarkably strong association between expression and patient survival for one gene that we had not previously examined. Low grade glioma (LGG) patients with high mRNA expression of the GAL3ST4 gene showed accelerated disease progression and poor survival when compared to patients with low expression. We also observed a similar trend when we examined copy number variations, where amplification of the GAL3ST4 gene was associated with poor prognosis (Fig. 6B). Finally, we found that methylation of the GAL3ST4 promoter region also predicted better patient survival (Fig. 6C). These data thus provided a strong impetus to investigate GAL3ST4 as a possible driver of immune evasion in glioma.

**Figure 6.**
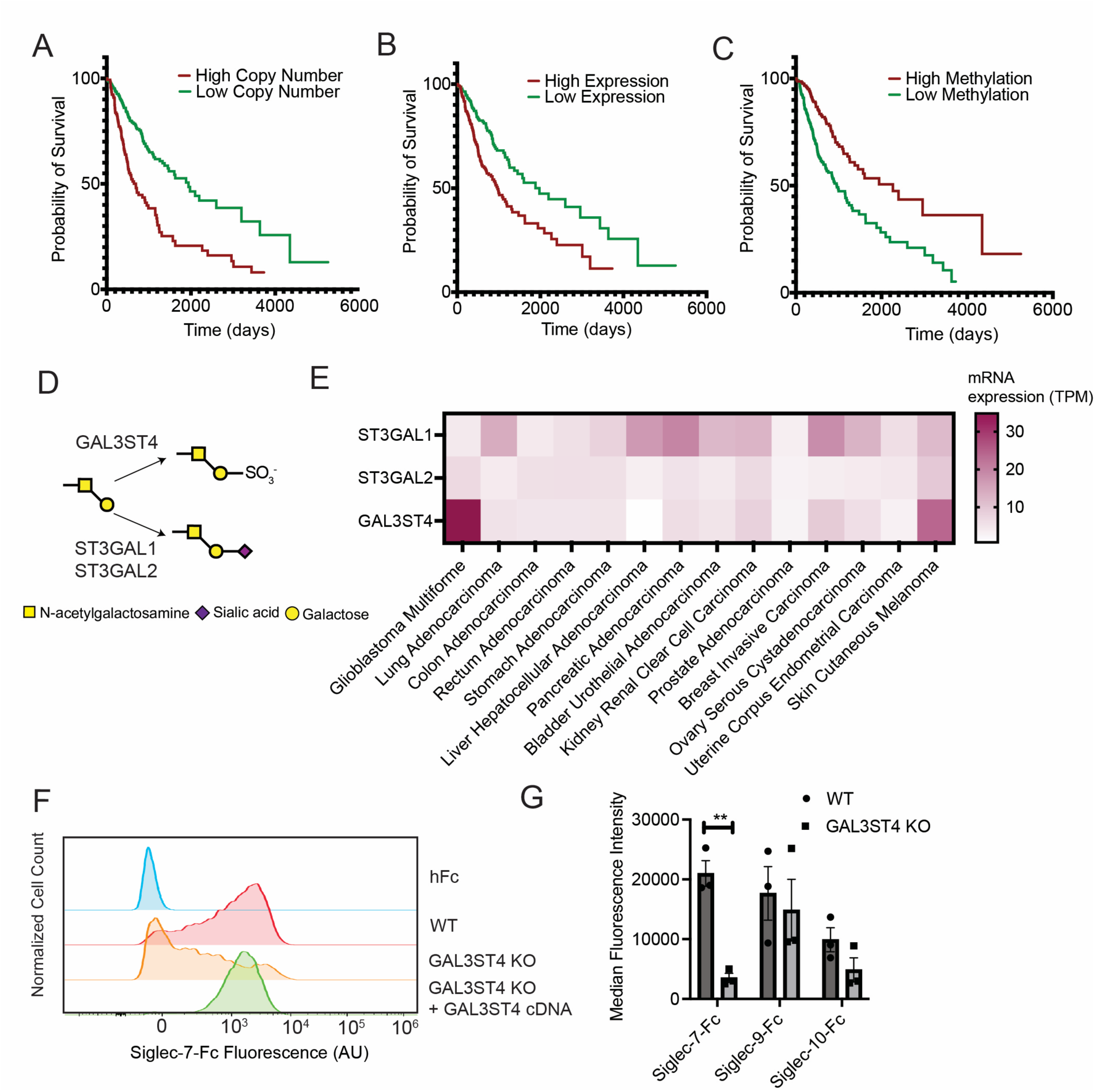
GAL3ST4 drives Siglec-7 ligand expression in glioma cells. **A)** Kaplan-Meier survival analysis was performed for low grade glioma (LGG) patients with either high or low chromosomal copy number at the GAL3ST4 locus. **B)** Kaplan-Meier survival analysis was performed for low grade glioma (LGG) patients with either high or low mRNA expression of the GAL3ST4 gene. **C)** Kaplan-Meir survival analysis was performed for low grade glioma (LGG) patients with either high or low DNA methylation at the GAL3ST4 promoter. Please see *Material & Methods* for details on bioinformatic analysis. **D)** Biosynthetic pathway for either sialylation (ST3GAL1/2) or sulfation (GAL3ST4) of core 1 O-linked glycans. **E)** The heat map indicates the average mRNA expression of ST3GAL1, ST3GAL2 and GAL3ST4 across 14 different cancer types. mRNA expression values were extracted from Human Protein Atlas. **F)** LN-229 cells were lentivirally transduced with Cas9 and an sgRNA targeting GAL3ST4. Cells were subsequently re-transduced with WT GAL3ST4 cDNA. LN-229 WT, GAL3ST4 KO and GAL3ST4 “rescue” cells were stained with fluorescent Siglec-7-Fc as described in **1B**. A representative flow cytometry plot is shown. **G)** LN-229 WT and GAL3ST4 KO cells were stained with fluorescent Siglec-Fc reagents as described in **1B**. The graph indicates the MFI for the three indicated Siglec-Fc reagents in WT and GAL3ST4 KO cells. Mean values plotted, ** indicates p<0.01, * indicates p<0.05. Mean values plotted, error bars indicate SEM.

Several recent studies demonstrated that Siglecs (particularly Siglec-9) are strongly upregulated on tumor-infiltrating myeloid cells in glioma and glioblastoma^15,81^. Inhibiting murine Siglecs also significantly reduced cancer progression in preclinical models of this disease^15,81^. However, the genetic perturbations that drive recruitment of Siglec-expressing immune cells have not been well-defined. To assess the role of GAL3ST4 in this process, we analyzed TCGA data to assess whether GAL3ST4 expression correlated with infiltration of Siglec-expressing immune cells in LGG. We compared mRNA expression of GAL3ST4 with that of Siglec-7 and Siglec-9 in a cohort of LGG patient samples. As Siglecs typically exhibit immune-restricted expression, we assumed that any mRNA expression of these Siglecs in a tumor sample would be derived from infiltrating immune cells and not tumor cells^11^. We found a strong correlation between GAL3ST4 expression and Siglec expression in these glioma patients (Supplementary Fig. 5). Our data thus imply a specific link between GAL3ST4, Siglec ligand expression and immune evasion in LGG.

GAL3ST4 is a sulfotransferase that appends sulfate to the 3’ hydroxyl group of galactose in O-linked glycans containing Galβ1,3-GalNAc (the T antigen)^82,83^. This gene emerged as a particularly strong hit in our Siglec-7 screen, where it was the top ranked positive regulator of Siglec-7-Fc binding. The finding that overexpression of sulfotransferases can drive Siglec ligand expression is supported by several prior studies^29,84^. The specific physiological context in which glycan sulfation is relevant for Siglec binding, however, has not yet been established. Sulfation of the T antigen by GAL3ST4 likely competes with sialylation of the same structure by ST3GAL1 and ST3GAL2 (Fig. 6D). In cells where ST3GAL1/2 expression is high, sulfation of the epitope is likely to be disfavored compared to sialylation. We therefore used the Human Protein Atlas to examine the average mRNA expression of GAL3ST4, ST3GAL1 and ST3GAL2 in 14 different cancers. In most cancer types, GAL3ST4 expression was very low (average TPM<5). ST3GAL1 and/or ST3GAL2 expression exceeded that of GAL3ST4 in almost all cases, indicating that sialylated core 1 structures are likely the dominant Siglec-7 ligands in most cases. Interestingly, the only strong exception to this trend was in glioblastoma, where GAL3ST4 expression significantly surpassed that of ST3GAL1/2. These data imply that GAL3ST4 may be a key tissue-specific driver of Siglec-7 ligand expression in brain cancer cells.

To test this hypothesis, we used the Human Protein Atlas to select an LGG cell line model (LN-229) with representative expression of GAL3ST4. We subsequently transduced this cell line with Cas9 and a sgRNA targeting GAL3ST4. Successful editing at the GAL3ST4 locus was confirmed by TIDE analysis (Supplementary Fig. 6). We then stained WT and GAL3ST4 KO cells with our Siglec-Fc chimeras. We observed a striking reduction in Siglec-7 ligand expression, with GAL3ST4 KO reducing Siglec-7-Fc binding to ∼20% of WT levels (Fig. 6F). Re-transfection of these cells with WT GAL3ST4 restored Siglec-7-Fc binding to WT levels, confirming the specificity of this effect. To our knowledge, this represents the first demonstration that inhibition of carbohydrate sulfation can ablate Siglec binding in a cancer cell model. GAL3ST4 could thus be a high-value target of interest for glioma and glioblastoma.

Interestingly, GAL3ST4 KO did not impact Siglec-9-Fc or Siglec-10-Fc binding in glioma cells (Fig. 6G). This finding reinforces that ligands for these different receptors can be generated by completely orthogonal driver genes. LN-229 cells, for example, exhibit quite high expression of genes like ST3GAL4, which is a key driver of Siglec-9 and Siglec-10 ligand expression (Human Protein Atlas). Several recent preclinical studies have highlighted the promise of Siglec-blocking antibodies as potential therapeutics for glioma and glioblastoma^15,81^. Our findings highlight the need to carefully consider the genetic background of a cancer when designing Siglec-blocking immunotherapy strategies. Siglec-7 and Siglec-9 induce redundant signaling pathways in immune cells. Siglec-9 blocking antibodies may thus be less effective in cases where tumor-intrinsic GAL3ST4 expression is high. Likewise, broadly acting therapeutics like sialidases may not effectively promote anticancer immunity in tumors where Siglec ligand expression is driven by carbohydrate sulfation^85^. Our findings thus urge closer integration of cancer genetic analysis with the design of immunotherapy regimens that target glycan-binding receptors.

## Discussion

We have applied a CRISPRa screening strategy to reveal key novel regulators of cancer glycome remodeling. One general finding was that expression of Siglec ligands is controlled by genetic circuits involving competing glycan-active enzymes. A key implication of this model is that there is no single gene whose dysregulation drives glycome remodeling across all cancers. The specific perturbation(s) required to achieve overexpression of Siglec-binding glycans will depend on the underlying transcriptional state of the healthy tissue. There are many genetic “paths” by which cancers may acquire a hypersialylated (or hypersulfated) phenotype, and these are likely to differ between tumor types and patients. In many cases, upregulation of specific STs may drive increased Siglec ligand expression. This has recently been demonstrated in several clinical contexts^13,17^. However, silencing of key glycan branching enzymes may be an equally potent mechanism for enhancing Siglec ligand expression in other tissues. Understanding these complex dynamics is becoming more important, as Siglec-blocking therapeutics are now being tested in clinical trials^15,41,42,86,87^. Currently, there are few reliable methods for identifying patients who are most likely to benefit from these therapeutics. Our CRISPRa dataset provides a multitude of possible biomarkers. In future, our work could be used to guide development of predictive gene expression models that can rapidly identify cancer patients likely to overexpress specific Siglec ligands.

Our screens also showed that Siglec binding is regulated by expression of key “professional ligands” that present glycans on the cell-surface. These genes are all cell-surface localized and thus accessible to biologic blocking therapeutics. This makes them high-value potential targets for cancer immunotherapy. Additionally, one strength of our GOF screening approach is that it has identified cell-surface targets that are relevant across a broad range of cancer types. For example, our screening data confirms that CD43 is likely the predominant Siglec-7 ligand in most types of blood cancer, while also revealing genes like MUC4 and MUC22 as potential targets in solid tumors. Likewise, our Siglec-9 screen revealed several heavily N-glycosylated and/or CS-modified proteins that are likely to serve as key ligands in different cell types (e.g., CSPG4 for melanoma)^73^. Our dataset thus provides a wealth of targets for future characterization in specific disease models. The biochemical features of these ligands that mediate Siglec binding will also be an interesting topic for future study.

Another key finding from our dataset is that expression of the cell-surface protein CD24 is not required for Siglec-10-Fc binding. There is an extensive literature indicating a connection between these two proteins, and our results in no way invalidate that prior work, which was largely conducted in quite different cells and model systems^41,75^. However, we do think our results urge a more precise examination of Siglec-10’s specificity for different types of glycoprotein ligands. Our data suggests that Siglec-10 is a receptor with a strong affinity for N-linked sialoglycans bearing both -2,3 and -2,6-linked sialic acid. Siglec-10 is thus likely to bind a range of cancer cell types with varying underlying patterns of GT expression. Unlike Siglec-7 and Siglec-9, we find no evidence of any additional specificity for specific glycoproteins. However, CRISPR screening is not a perfect method, and this topic deserves a more focused examination in future work.

Finally, our work identifies GAL3ST4 as a possible driver of immune evasion in glioma. This enzyme has several appealing properties as a potential drug target. Firstly, the gene is selectively upregulated in glioma patient samples and is a clear adverse prognostic factor. This contrasts with other glycan-active enzymes, where we found it difficult to find prognostic associations in clinical data. Secondly, expression of this enzyme is quite tissue-restricted relative to the other glycan-active enzymes (ST3GAL1, ST3GAL2, ST6GALNAC4, etc.) that generate Siglec-7 ligands. In principle, pharmacological inhibition of this enzyme may have limited off-target effects. While small molecule sulfotransferase inhibitors have been explored as drug candidates, there are currently no known specific inhibitors of GAL3ST4^88^. If these cannot be developed, our data also imply that blocking Siglec-7 may be an effective strategy for attacking gliomas with high expression of GAL3ST4. We plan to explore these possibilities in future work.

Taken together, these studies demonstrate the power of GOF CRISPRa screening for dissecting regulation of glycan biosynthesis pathways. The continued development and optimization of our FACS-based screening platforms create new opportunities for study of other glycan-binding receptors in different disease contexts. In future, we view CRISPRa screening as a valuable tool for comprehensively mapping the genetic factors that drive glycosylation in living cells.

## Acknowledgements

Prof. Wisnovsky acknowledges funding support from the National Science & Engineering Research Council of Canada (NSERC), the Canadian Institutes of Health Research (CIHR), the Canadian Glycomics Network, the Cancer Research Society, the Canadian Cancer Society and Michael Smith Health Research BC.

## Materials & Methods

### Cell culture

All cell lines used in this study were acquired from American Type Culture Collection (ATCC). K562, THP-1 and MM1S were cultured in Roswell Park Memorial Institute 1640 (RPMI-1640) medium supplemented with 10% fetal bovine serum (FBS, Sigma) at 5% CO2 and 37 °C. OCI-AML2 and Lenti-X HEK293T cells were cultured in Dulbecco’s Modified Eagle Medium (DMEM) supplemented with 10FBS at 5% CO_2_ and 37°C. LN-229 cells were cultured in DMEM media containing 5% FBS at 5% CO2 and 37°C. All cell lines were sub-cultured at frequencies and concentrations recommended by the supplier.

### Siglec-Fc staining

Fluorescently labelled Siglec-Fc chimeras were prepared by pre-complexing 1.5µg/mL of Siglec-7Fc, Siglec-9Fc and Siglec-10Fc chimeras (R&D Biosystems: 1138-SL, 1139-SL and 2130-SL, respectively) with 1.5µg/mL Alexa Fluor® 647 AffiniPure™ Goat Anti-Human IgG, F(ab’)₂ fragment specific, (Jackson ImmunoResearch: 109-605-006) in 1X PBS for 45mins on ice in the dark. As a control 1.5µg/mL Recombinant Human IgG1 Fc (Thr106-Lys330), (BioLegend: 773006) was pre-complexed with Alexa Fluor® 647 AffiniPure™ Goat Anti-Human IgG, F(ab’)₂ fragment specific under the same conditions. Target cells were often co-stained with fluorescently labelled Siglec-Fc chimeras and lectins. In this case, L-PHA (5µg/mL) or DBA (10µg/mL) were added to Siglec-Fc precomplexes during incubation. Cells were stained on ice in the dark for 30mins, before being centrifuged at 600xG for 5mins after which supernatant was discarded, cells were washed with 1X PBS, resuspended in 1X PBS and samples were run on a BD LSR II (BD Biosciences) flow cytometer. Meanwhile, cells of interest were collected, counted, washed with 1X PBS and aliquoted into 96-well V-bottom culture plates (Corning Incorporated) before being centrifuged at 600xG for 5mins. Supernatant was discarded and cells were then resuspended in appropriate pre-complex solution at 1x10^5^ cells/100µL (for K562, THP-1, MM1S and LN-229), while 2x10^5^ cells/100µL was used for OCI-AML-2. Cells were incubated for 30 min on ice in 96-well V-bottom plates. Cells were subsequently centrifuged at 600xG for 5mins after which supernatant was discarded, cells were washed with 1X PBS, resuspended in 1X PBS and samples were run on either an LSR II (BD Biosciences) or a CytoFLEX (Beckman Coulter) flow cytometer. FSC vs SSC was used to gate intact cells and FSC-A vs FSC-H was used to exclude doublets. For cells transduced with virus targeting a single gene (for either gain-of-function or knockout), cells were further gated to include solely BFP^+^ cells before expression of the antigen of interest was recorded. A minimum of 5,000 single, intact cells were recorded for each sample.

### CRISPR activation (CRISPRa) screening

#### Packaging of Genome-Wide CRISPRa Library

The CRISPRa-v2 library (top 5 sgRNAs/gene) containing 104,535 sgRNAs was purchased from Addgene (Pooled Library 83978). For lentiviral packaging, HEK293T cells were plated onto six 150 mm dishes at a cell concentration of 7.5x10^6^ cells/dish in 30mL of medium and allowed to adhere overnight. For each plate, 8 µg of sgRNA library plasmid was mixed with 8 µg total lentiviral packaging plasmids (4µg PMD2.G and 4µg psPAX2) in 2mL of serum-free DMEM. 48 µL of LT1 (Mirus Bioscience) transfection reagent was separately diluted in 2mL of serum-free DMEM and allowed to sit at room temperature for 5mins. The two solutions were then combined to create a transfection complex and the mixture was allowed to incubate at room temperature for 30mins. The total volume of each transfection complex was then carefully added to each 150mm plate dropwise. During this process the media on the plate was swirled slowly but constantly. Cells were then placed back in the incubator and virus production was allowed to proceed for 72hrs. Lentiviral media was then removed from each plate, centrifuged at 3000xg for 10mins to clear any cellular debris. Lentiviral media was then snap frozen and stored at -80°C until the day of library infection.

#### Lentiviral Transduction of Genome-Wide CRISPRa Library

2.5x10^8^ K562-dCas9-VP64 cells growing in log phase were spun down and resuspended in 500mL of complete media containing 8µg/mL polybrene and a volume of lentiviral media previously determined to give a multiplicity of infection (MOI) of 0.35. MOI was quantitated in a smaller scale experiment by determining the titer of lentiviral media that produced 30% BFP expression (as determined by flow cytometry) 72hrs following infection. Cells were then selected with 1µg/mL of puromycin for 48hrs. Cells were then aliquoted in 10 separate 150 mm plates (50mL per plate). After 24hrs, cells were centrifuged at 600xG and resuspended in media containing 1 µg/mL of puromycin at 5x10^5^ cells/mL. Following 48hrs of puromycin selection, cells were spun down and resuspended again in fresh media containing 1 µg/mL of puromycin, maintaining a cell density of 5x10^5^ cells/mL. After 96hrs of selection, viability of the infected cell population was greater than 95%, while an uninfected control plate similarly treated with puromycin had cellular viability less than 1%. After this point, cells were spun down and resuspended in fresh media without puromycin and expanded. Cell staining and sorting in all cases were performed within 5 days of removing cells from puromycin selection to avoid dropout of essential genes from extended maintenance of the library in culture.

#### Siglec Sorting of Genome-Wide CRISPRa Libraries

Siglec-7, Siglec-9 and Siglec-10 CRISPRa screens were performed in duplicate. At all points, all replicates were maintained in culture at a library coverage of 1000x (∼1x10^8^ cells/replicate). Prior to sorting, 2µg/mL of Siglec-7Fc, Siglec-9-Fc or Siglec-10Fc was pre-complexed with 2µg/mL of Alexa Fluor® 647 AffiniPure™ Goat Anti-Human IgG, F(ab’)₂ fragment specific in a total volume of 24mL of 1X PBS buffer in the usual manner and incubated on ice for 45mins to precomplex. For each CRISPRa genome-wide screen replicate, 1.2x10^8^ cells were then pelleted at 600xG, washed once with 1X PBS and resuspended in precomplex solutions at 5x10^6^ cells/mL. Cells were incubated on a rocker table in a cold room for 45mins to ensure consistent but gentle mixing of the cells. Following staining, cells were spun down at 600xG, washed twice with 1X PBS and resuspended in 1X PBS. Cells were then passed through a 70µm nylon cell strainer (Corning Incorporated) to remove any aggregates and then placed on the sorter arm, where they were kept at 4 °C and rotated frequently for the duration of the sort. Cell sorting was performed on a BD FACSAria II. Intact BFP^+^, single cells were selected by gating on FSC vs SSC (to exclude debris), FSC-A vs FSC-H (to exclude doublets) and FSC-H vs BV421 (to exclude BFP^-^ cells) parameters. Finally, gates on Siglec-Fc staining were constructed such that cells were only sorted if they exhibited a fluorescence intensity value less or more than 10-fold than the average fluorescence value of the stained population. In practice, this usually corresponded to approximately 40% of the total stained population (20% of the lowest and 20% of the highest fractions as determined by mean fluorescence intensity depending on the specific Siglec-Fc fluorescence distribution). Sorted cells were periodically pelleted and genomic DNA was immediately extracted before being quantified and frozen. Sorting was performed until the whole input cell sample had been totally depleted.

#### Library Amplification, Sequencing & Data Processing

DNA from sorted samples was isolated using the Sigma GeneElute Genomic DNA Miniprep Kit according to manufacturer’s instructions. Libraries were amplified via nested PCR and sequenced on an Illumina NextSeq as previously described^26^. Following demultiplexing, FASTQ sequence alignment to library file was conducted using guide counter V0.1.3. sgRNA counts from the two low-staining replicates were compared to sgRNA counts from two high-staining replicates using CasTLE (**Cas**9 High **T**hroughput maximum **L**ikelihood **E**stimator)^44^.

### Bioinformatic Analysis

#### CRISPR Screen Analysis & Hit Identification

FASTQ files from each sample were aligned to the reference CRISPRa library using guide-counter to generate a read count table. Read count tables were then analyzed using CasTLE^44^ to identify sgRNAs that exhibited altered abundance in low-staining and high-staining samples. If sgRNAs targeting a given gene were more abundant in the high-staining sample than the low-staining sample, the gene was classified as a hit with a positive effect score. If sgRNAs targeting a given gene were more abundant in the low-staining sample than the high-staining sample, the gene was classified as a hit with a positive effect score.

#### Gene Ontology (GO) Enrichment Analysis

Enriched GO terms in each screen were identified by ranking all gene hits by CasTLE Score (both positive and negative hits). This ranked list was then analyzed using GORilla^89^. A false discovery rate cutoff of 0.01 was used to identify statistically significant GO terms. This filtered list was then ranked by fold enrichment prior to displaying data.

#### TCGA Analysis

Genome-wide survival scores for all top screen hits were accessed using a previously published resource^80^ (https://www.tcga-survival.com/). In this survival model, all genes are assigned a Z-score in each cancer type. A higher Z-score indicating that genetic variation in that gene has prognostic significance. Gene copy number, methylation and mRNA expression were all analyzed separately. Genes with a Z-score above 5 in at least one cancer type were prioritized.

#### mRNA expression analysis

mRNA expression data for human cell lines was sourced from the Human Protein Atlas (proteinatlas.org). mRNA expression data for clinical samples was sourced from the TCGA (cbioportal.org) using the Brain Lower Grade Glioma (TCGA, PanCancer Atlas) dataset.

### Lentiviral transduction of sgRNAs

Lentivirus was prepared as previously described^13^. Lentiviral supernatant was collected and spun for 10mins at 1,000xG to pellet any floating cells. Supernatant was carefully removed to avoid disturbing the pellet, snap frozen in 500µL aliquots and stored at -80°C until day of use. On the day of transduction, cells from relevant lines were seeded at recommended subculture concentrations in appropriate culture media for a total volume of 2mL in a 6-well cell culture plate (Corning). In total, two wells were seeded to include a control for selection. 500µL of viral supernatant (or standard culture media as a control) was added to the cells along with 8µg/mL of polybrene (MilliporeSigma: TR-1003-G). The culture was gently mixed and incubated at 37°C for 48hrs after which the cells were collected (at this stage an aliquot of cells was collected and examined for BFP expression using flow cytometry to determine the initial viral transduction efficiency) and centrifuged for 5mins at 600xG. Supernatant was removed and cells were resuspended in 3mL of appropriate culture media containing 1µg/mL puromycin (InvivoGen: ant-pr) before incubation at 37°C for 48hrs. Media was replaced every 48hrs along with 1µg/mL puromycin until all there were no viable cells in the control well. If needed, puromycin concentration was increased. After selection, cells were cultured at recommended densities until aliquots were frozen or used in phenotyping assays (flow cytometry, TIDE analysis).

For MM1S cells, virus was prepared and stored in the usual manner^13^. RetroNectin-coated plates (Takara: T100A/B) were used to enhance transduction efficiency. Supplier’s instructions were followed to ensure successful coating of the plates with virus and transduction of target cell lines. Minor modifications were made to the supplier’s protocol depending on the sub-culture seeding densities and doubling time of specific cell lines. After incubation with lentivirus (between 48-96hrs, depending on the cell line used), an aliquot of cells was collected and examined for BFP expression using flow cytometry to determine initial transduction efficiency. ST6GAL1 KO cells were then isolated via lectin staining and FACS. 3x10^7^ MM1S ST6GAL1 KO cells were stained with 13.33µg/mL of *Sambucus Nigra* Lectin (SNA, Invitrogen: L32479) in a final volume of 15mL (2x10^6^ cells/mL) for 30mins on ice in the dark. Cells were washed with 1X PBS and then sorted using a BD FACS Aria III (BD Biosciences). Briefly, Forward Scatter vs Side Scatter (FSC vs SSC) gating was used to exclude debris and FSC-Area vs FSC-Height (FSC-A vs FSC-H) gating was using to exclude doublets. Cells were then gated to include solely BFP^+^ cells after which cells were examined for SNA binding. As SNA binds to α2,6-linked sialic acids, SNA^+^ and SNA^-^ cells were present. Using the BD FACSAria II (BD Biosciences), SNA^-^ cells were sterile sorted, counted, washed and re-suspended in culture medium at 5% CO_2_ and 37°C. These cells were cultured until aliquots were frozen or until needed in downstream phenotyping assays.

### Genomic DNA PCR & TIDE analysis

Genomic DNA was extracted using the GeneJET Genomic DNA Purification Kit (ThermoFisher, K0721) according to the manufacturer’s protocol. Fragments containing the sgRNA target sites were subsequently amplified via PCR with the Herculase II Fusion DNA Polymerase kit (Agilent, 600675). The resulting PCR products were resolved on a 1% agarose gel, purified using the GeneJET Gel Extraction Kit (ThermoFisher, K0691), and submitted to the UBC Sequencing and Bioinformatics Consortium for Sanger sequencing. Sequence data from wild-type and knock-out samples were analyzed with TIDE software, which quantifies insertion and deletion events at the target locus through deconvolution of Sanger sequencing traces.

**Table 1.** Key Antibodies & Recombinant Proteins Used in Study.

| Reagent | Vendor | Catalogue Number |
| --- | --- | --- |
| Siglec-7-Fc | R&D Systems | 1138-SL |
| Siglec-9-Fc | R&D Systems | 1139-SL |
| Siglec-10-Fc | R&D Systems | 2130-SL |
| Recombinant Human IgG1 Fc (Thr106-Lys330) | BioLegend | 773006 |
| Alexa Fluor® 647 AffiniPure™ Goat Anti-Human IgG | Jackson Immuno Research | 109-605-008 |
| PE-anti-human-CD24 (Clone W20001B) | BioLegend | 382609 |
| PE-Mouse IgG2b,k Isotype Control | BioLegend | 402203 |

**Table 2.**
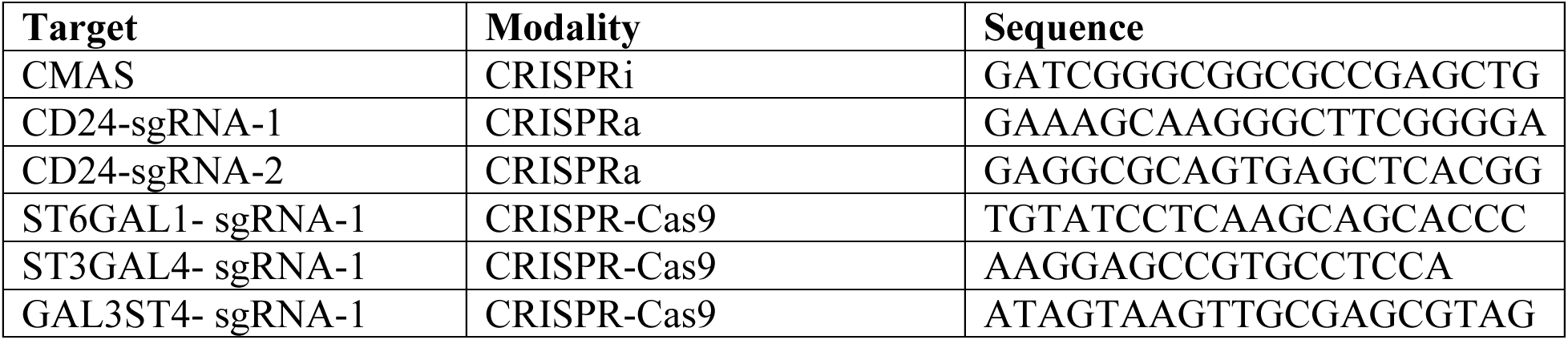
sgRNA Sequences Used for CRISPR Engineering.

**Table 3.**
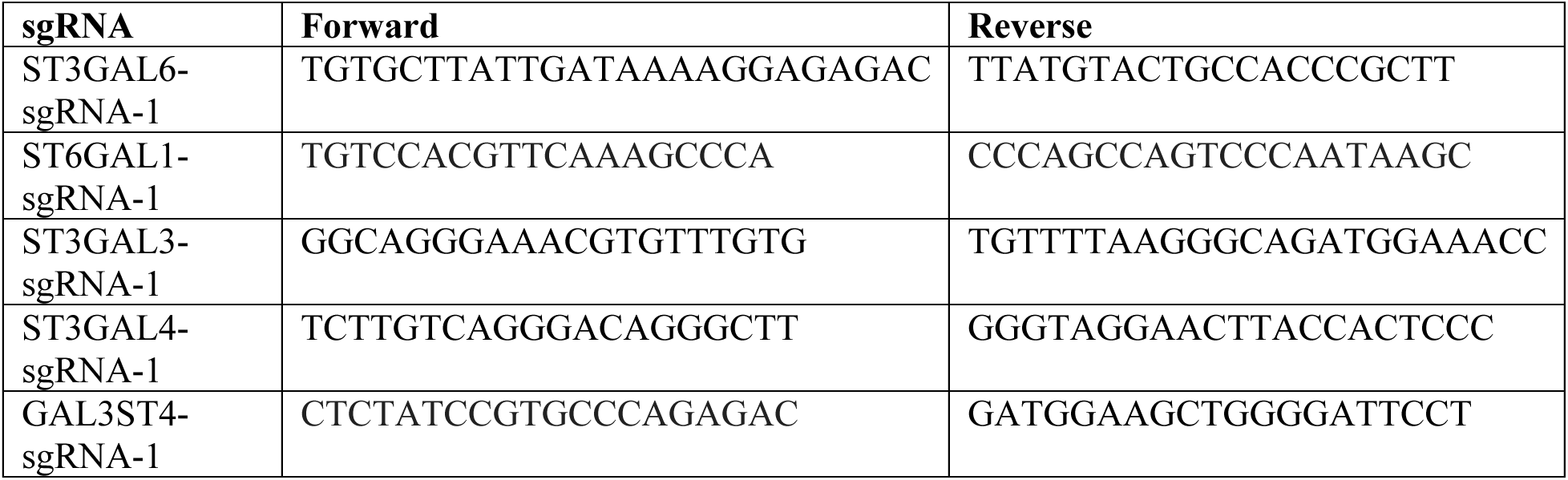
Primer Sequences Used for TIDE Analysis.

