## Supplementary Information for "CRISPR activation screens map the genomic landscape of cancer glycome remodeling"

­­

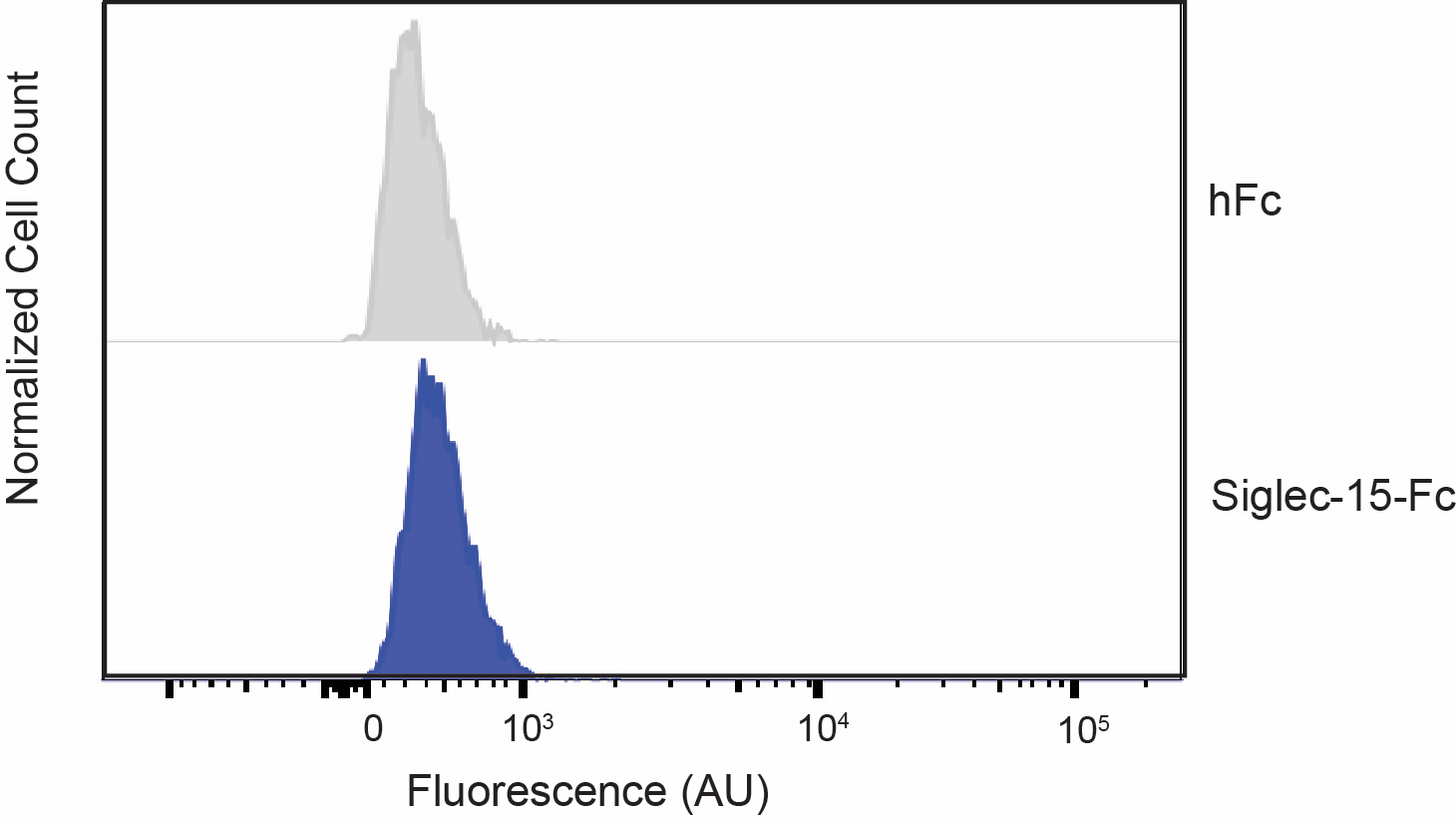

**Supplementary Figure 1.** K-562-CRISPRa cells were incubated with Siglec-15-Fc precomplexed to a fluorescent secondary antibody. A representative flow cytometry plot is shown.

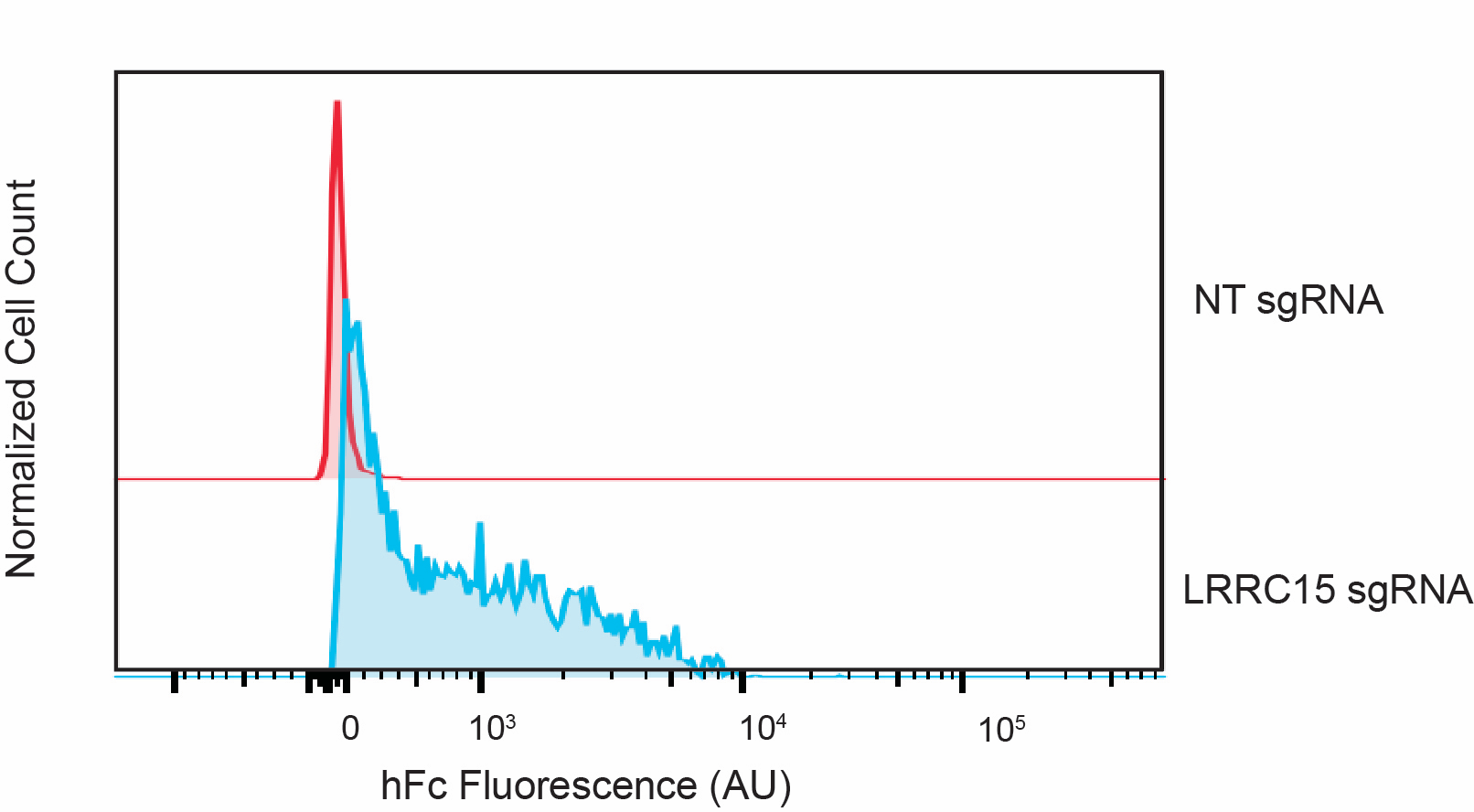

**Supplementary Figure 2.** Transduction of K-562 CRISPRa cells with an sgRNA targeting LRRC15 leads to increased hFc binding.

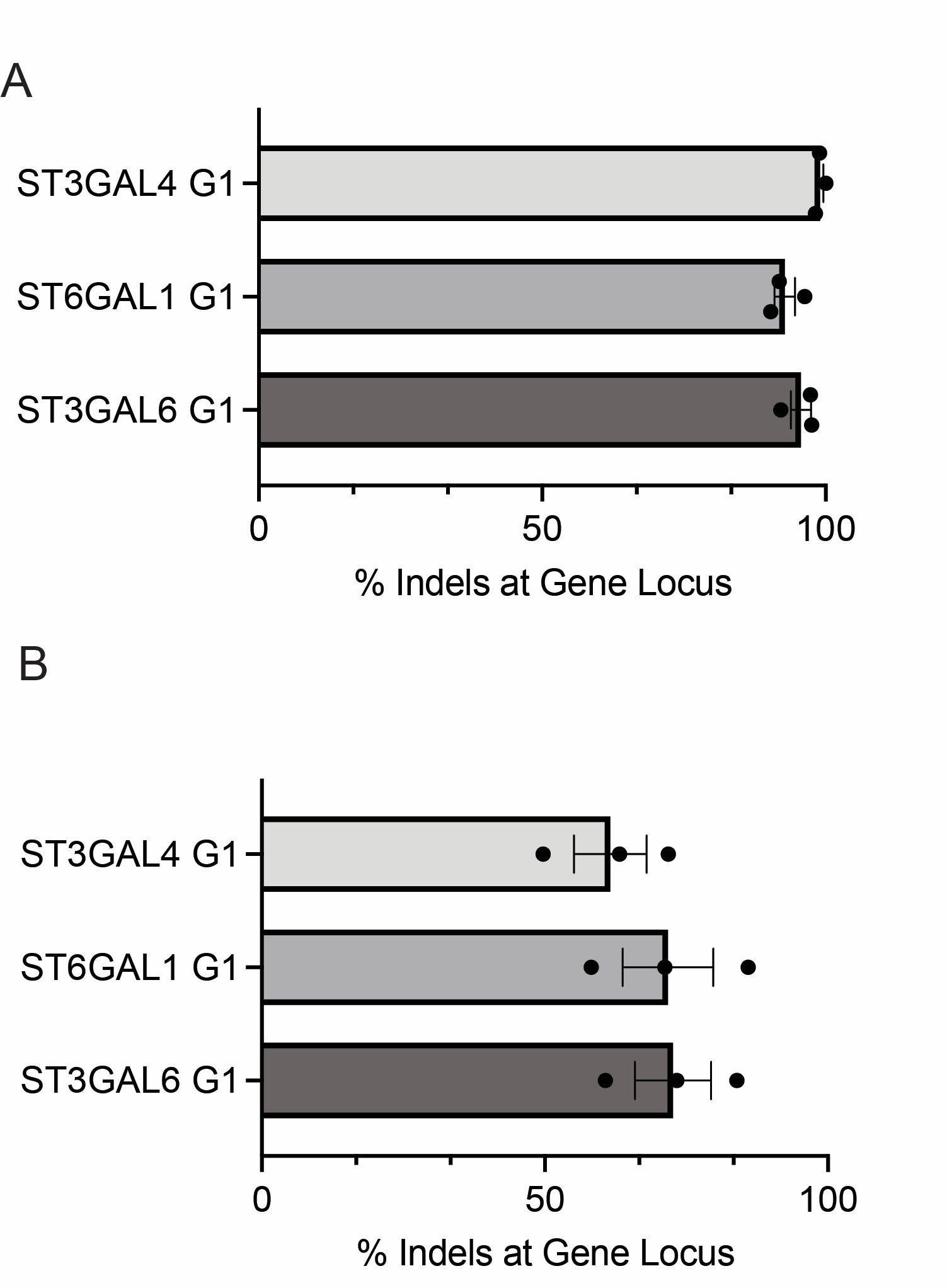

**Supplementary Figure 3.** TIDE analysis of gene knockouts in OCI-AML-2 cells. Error bars indicate SEM, n=3.

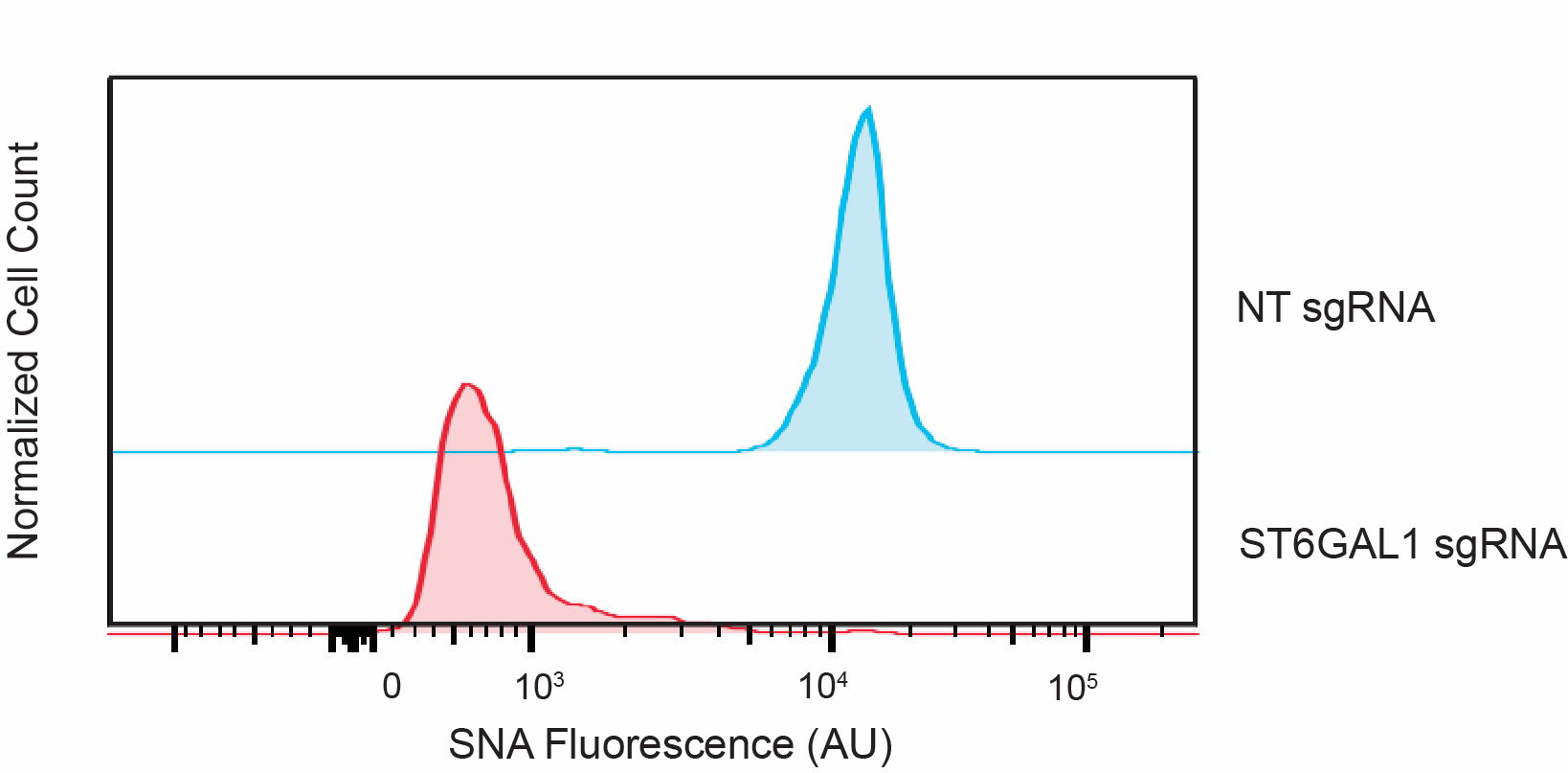

**Supplementary Figure 4.** MM1S-Cas9 cells were transduced with an sgRNA targeting ST6GAL1, stained with fluorescent SNA and sorted via FACS to isolate cells with decreased expression of 2,6-linked sialic acids.

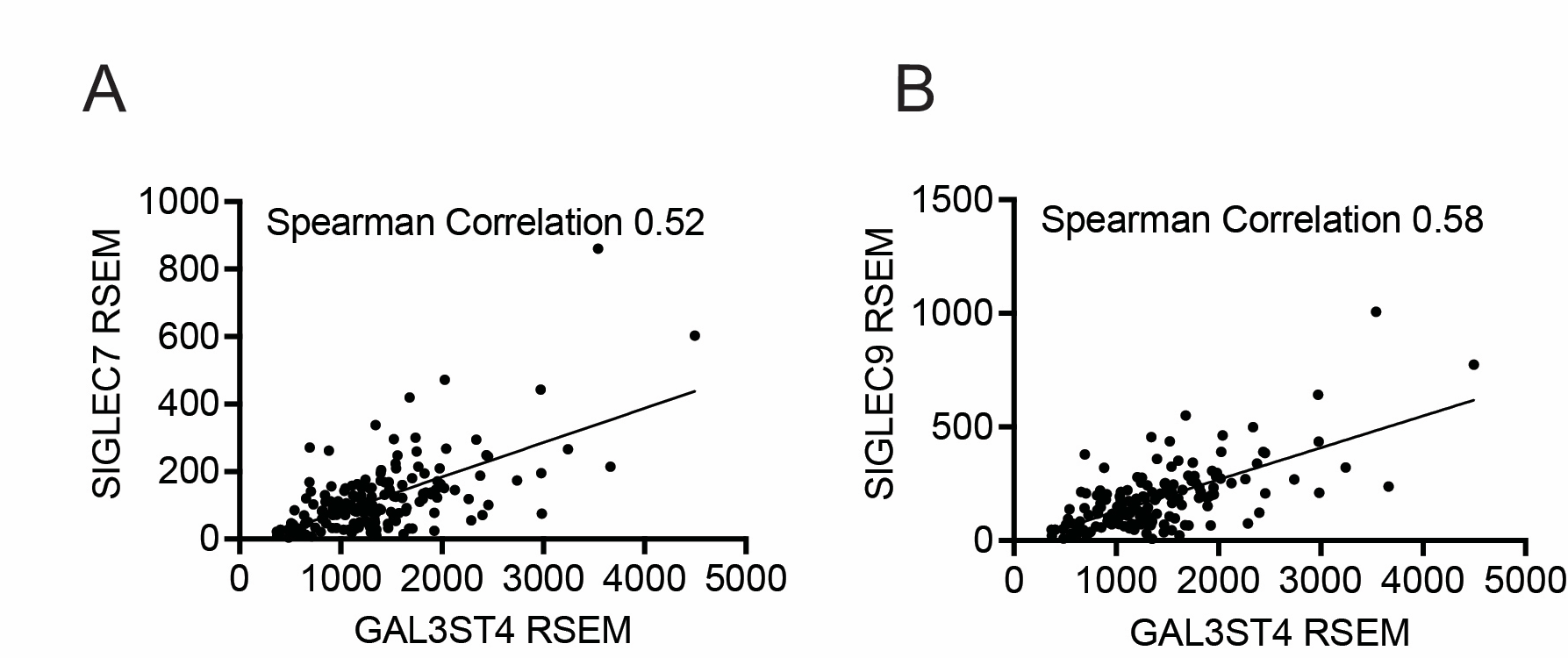

**Supplementary Figure 5. A)** The graph indicates the correlation between mRNA expression of GAL3ST4 and Siglec-7 in primary low grade glioma (LGG) patient samples. **B)** The graph indicates the correlation between mRNA expression of GAL3ST4 and Siglec-9 in primary low grade glioma (LGG) patient samples. mRNA expression data was extracted from TCGA as described in *Material & Methods.*

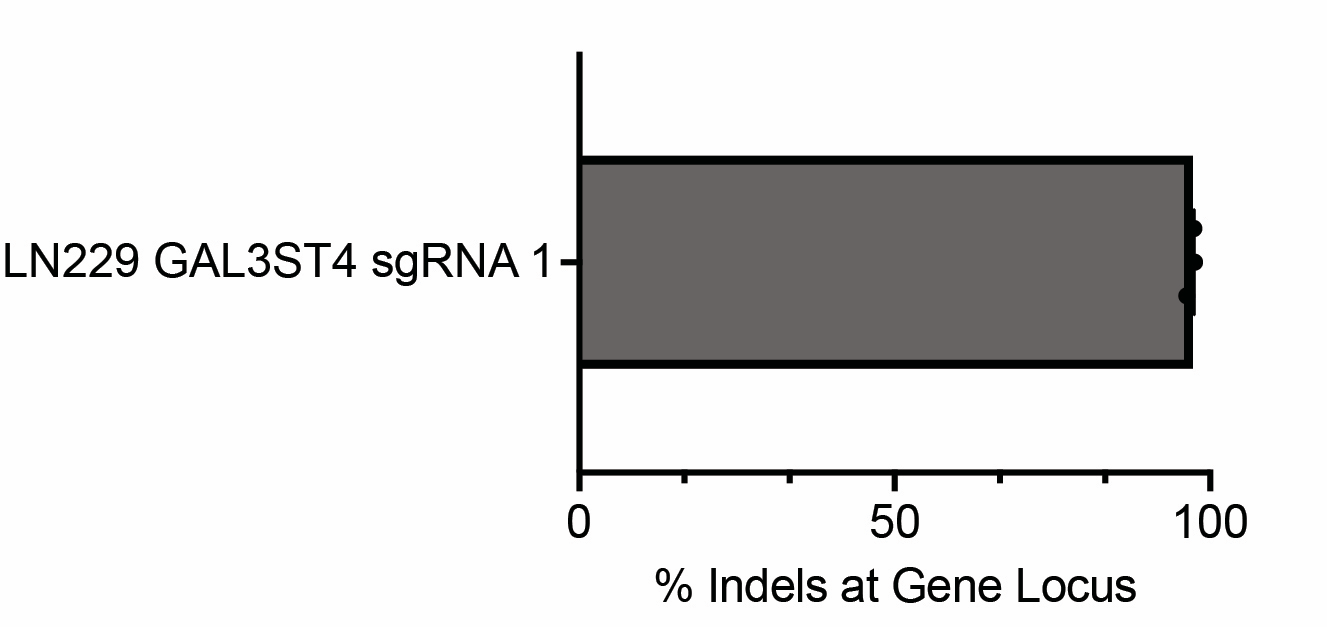

**Supplementary Figure 5.** TIDE analysis of GAL3ST4 KO in LN-229 glioma cells. Error bars indicate SEM, n=3.
